# Spatial entropy of brain network landscapes: a novel method to assess spatial disorder in brain networks

**DOI:** 10.1101/2025.10.23.684130

**Authors:** Clayton C. McIntyre, Shannon M. O’Donnell, Mohammadreza Khodaei, Robert G. Lyday, Jonathan H. Burdette, Paul J. Laurienti

**Author notes:** **Corresponding Author:** Paul J. Laurienti, Department of Radiology, Wake Forest University School of Medicine, Medical Center Boulevard, Winston-Salem, NC 27157. These authors contributed equally (Clayton C. McIntyre, Shannon M. O’Donnell).

## Abstract

In this work, we introduce a method for mapping the spatial entropy of functional brain network community structure images in brain space. Entropy maps indicate the extent to which the network communities present in a local area are ordered or disordered. We demonstrate how spatial entropy can be quantified for each voxel in the brain according to the network community affiliations of surrounding voxels. This process results in interpretable maps of brain network entropy. We show that local entropy decreases in predictable brain regions during working memory and music-listening tasks. We suggest that these regional entropy reductions reflect self-organization of neural processes in support of functionally localized cognitive tasks. Analyses in this work provide a framework for future analyses of spatial entropy in complex networks that can be mapped to Euclidean space – both within the brain and in other contexts.

**Significance Statement:** We introduce an approach for quantifying the spatial entropy of functional brain network community structure. We demonstrate the biological relevance of the measure in three independent datasets. This approach for analyzing brain network data is data-driven, easy to implement, and highly interpretable. It also allows investigators to visualize complex data by mapping values into the brain rather than storing values in extremely high-dimensional and abstract data structures. We believe this will make the method highly accessible even to investigators with minimal experience analyzing human neuroimaging data.

## Introduction

The human brain is a complex system that relies on coordinated spatial organization of networks (Bullmore & Sporns, 2009). A fundamental principle of this organization is the balance between integration and segregation of information, which supports communication across brain regions and specialized processing within distinct subnetworks. Disruptions in this organization impedes the brain’s ability to manage information transfer and processing which can result in neurological or psychological disorders (Peng et al., 2014). Identifying patterns of spatial organization that govern these processes can give us valuable insight into understanding normal vs abnormal brain function. One way to quantify these spatial organization patterns is by partitioning brain networks into communities and examining their properties.

Community structure is determined solely based on the topology of the network without taking into account the physical location of the brain regions that nodes represent (Bertolero et al., 2015; Kabbara et al., 2019). However, once community assignments are identified for each node, they can be mapped into brain space. Conceptually, this should provide a very enlightening view of functional imaging data. In fact, communities have neurobiological relevance as their spatial distributions can reveal functionally specialized subsystems (Sporns & Betzel, 2016). Unfortunately, there are limited methods for analyzing spatial structure of network communities at the group level. This problem is multifaceted as there is not a “ground truth” community structure to align the functional brain network communities to. Moreover, these communities don’t align to the same location across participants. Methods have been developed in an attempt to address this problem but often include creating a representative community division (i.e., group average) for the groups (Kirkley & Newman, 2022) or by using a priori spatial definitions (Moussa et al., 2012; Steen et al., 2011) resulting in a potentially inaccurate depiction of the data. An ideal method would allow analysis of the spatial organization of brain network communities in a purely data-driven approach.

Information theory, developed by Claude Shannon, is a mathematical concept originally used to analyze data transmission for communication purposes (Shannon 1948). At the center of the theory is the idea that the entropy of a random variable is defined as the amount of “uncertainty”, “surprise”, or “diversity” in the possible outcomes of the variable (Ben-Gal & Kagan, 2021). In other words, a more disordered system has higher entropy (Saraiva, 2023). However, there are two major limitations when applying Shannon’s entropy to spatial systems. First, Shannon’s original equation only takes into consideration the probability distribution of the variable under study. For spatial applications, the focus is not purely on probability distributions but also involves the spatial distribution of variables in the system (Altieri et al., 2023). Second, space is multidimensional. In Shannon’s original work, the focus of study was one-dimensional signals (Wang & Zhao 2018). Due to these limitations, methods to quantify entropy for spatial problems have been adapted from Shannon’s equation.

A key step in the development of spatial entropy methods came from the ecology literature with the introduction of Batty’s entropy (Batty, 1974). This method divided one large area into many subareas by applying a grid. Entropy could then be calculated separately for subareas, which allows for spatial specificity and the ability to create a “map” of entropy. This approach can be applied to functional brain network community structure very easily because the data is collected in a 3-dimensional grid structure. Subareas are easily defined as spatial clusters of voxels (more generally, voxels can be considered the “elements” of the subarea). Because brain anatomy is practically identical across people, brain network entropy maps can be directly compared at a group level. While the precise implementation of spatial entropy as proposed by Batty is not compatible with brain network analyses, Batty’s proposal inspired creation of several subsequent measures of spatial entropy that do work for brain networks (O’Neill et al. 1988; Claramunt, 2005; Wang & Zhao, 2018). In this work, we investigate four spatial entropy measures.

The first measure (referred to as spatial Shannon’s entropy) utilizes subareas similar to Batty and calculates the Shannon entropy within each subarea based on the community probabilities. The second measure quantifies how communities are arranged within a subarea by focusing on probability of contiguous (touching) pairs of communities (O’Neill et al., 1988). The third measure incorporates distance within and between communities in each subarea (Claramunt, 2005). The fourth measure serves as a conceptual hybrid of the second and third measures by accounting for the probability of shared borders between communities as well as the distance between community centroids within the subarea (Wang & Zhao, 2018). Computational details of each entropy measure are described below.

The present study had two primary aims. The first was to apply each of the four entropy measures to simulated spatial patterns that were simple enough to have interpretable outcomes. Given the complexity of the brain, it can be difficult to understand how specific spatial patterns of community structure would be captured by the different entropy measures. We aimed to determine differences across all four entropy measures when calculating one entropy value for the whole system (as proposed in Batty’s work intended for ecological systems) as well as calculating an entropy value in every subarea to create an entropy map.

Our second aim was to apply the four entropy measures to real brain community structure landscapes derived from functional magnetic resonance imaging (fMRI) data to 1) characterize differences in entropy maps based on the different measures and multiple scaling factors, and 2) demonstrate that differences in entropy maps corresponding to changes in cognitive/sensory state have biological meaning. To do this, we sought to determine which regions of the brain displayed relatively higher or lower entropy levels during cognitive and auditory tasks as compared to resting-state. Previous work has shown that nodes of the default mode network (DMN) are tightly integrated during rest, whereas nodes of the central executive network (CEN) reorganize into a highly integrated community during working memory (Moraschi et al., 2020). As such, we hypothesized that the DMN would have relatively low entropy at rest while regions associated with the CEN would have relatively low entropy during a working memory task. We replicated our findings using an independent dataset with resting-state and working memory scans. We also assessed spatial entropy of community structure landscapes in a music-listening study. We hypothesized that entropy would decrease in the auditory cortex when participants were engaged with an auditory stimulus in the scanner.

### Methodology

#### Spatial Entropy Measures

We examined the spatial disorder of simulated systems and functional brain network community structure using four measures of spatial entropy. Spatial entropy was calculated for each and every element (i.e., voxel) based on the community assignments of the elements in the subarea of the central element (for detailed description of element/voxel subareas, see Detailed Methods). Calculating entropy for every element meant that the subareas attributed to adjacent voxels tended to have a substantial amount of overlap.

The first entropy measure (*H*_*S*_) is a spatial form of Shannon’s entropy (Shannon, 1948) that we adapted from Batty’s method (Batty, 1974) to accommodate the data structure of brain network community maps. It is calculated for each subarea of elements as

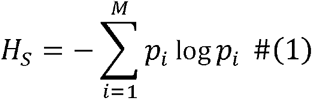

where *M* is the number of communities in the system and *p*_*i*_ is the proportion of elements in the subarea belonging to community *i*.

The second entropy measure (*H*_*O*_) was introduced by O’Neill (O’Neill et al., 1988). This measure is focused on the shared borders between adjacent elements within the subarea, and is calculated as

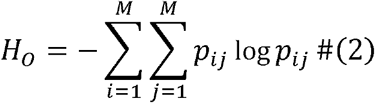

where *p*_*ij*_ is the proportion of shared edges between elements of communities *i* and *j*.

The third entropy measure (*H*_*C*_) was introduced by Claramunt (Claramunt, 2005). This measure adds a scaling factor to the *H*_*S*_ calculation to adjust for the distance between elements of similar and different communities within the subarea. It is calculated as

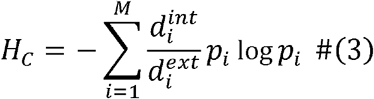

where 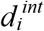 is the average distance between elements in community *i* and 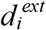 is the average distance between elements belonging to community *i* and elements belonging to other communities.

The final entropy measure, proximity entropy (*H*_*P*_), was introduced by Wang and Zhao (Wang & Zhao, 2018). This measure is conceptually a combination of *H*_*O*_ and *H*_*C*_. It is computed by adding a scaling factor to the *H*_*S*_ calculation based on the shared borders between communities and the distance between community centroids. It is calculated as

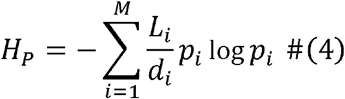

where *L*_*i*_ is the number of edges between elements of community *i* and elements of different communities and *d*_*i*_ is the sum of the distances between the Euclidean centers of community *i* and all other communities.

Scripts used to generate brain entropy maps based on community landscapes are openly available at https://github.com/rlyday/brain-network-spatial-entropy.

## Results

### Spatial Entropy in a Simulated System

Figure 1 demonstrates the difference between calculating a single entropy value for large, simulated spatial systems versus dividing them into distinct subareas and calculating spatial entropy within subareas. In Figure 1a, each entropy measure is used to calculate the system-level entropy (i.e., no parcellation into subareas) for two simulated spatial systems. While the system on the right is clearly more spatially disordered than the system on the left, *H*_*S*_ is equal between systems because the number of elements belonging to each community is constant across systems. The other three entropy measures (*H*_*O*_, *H*_*C*_, and *H*_*P*_) were developed to address this limitation to *H*_*S*_. As expected, each of these entropy measures is higher for the system on the right.

**Figure 1.**
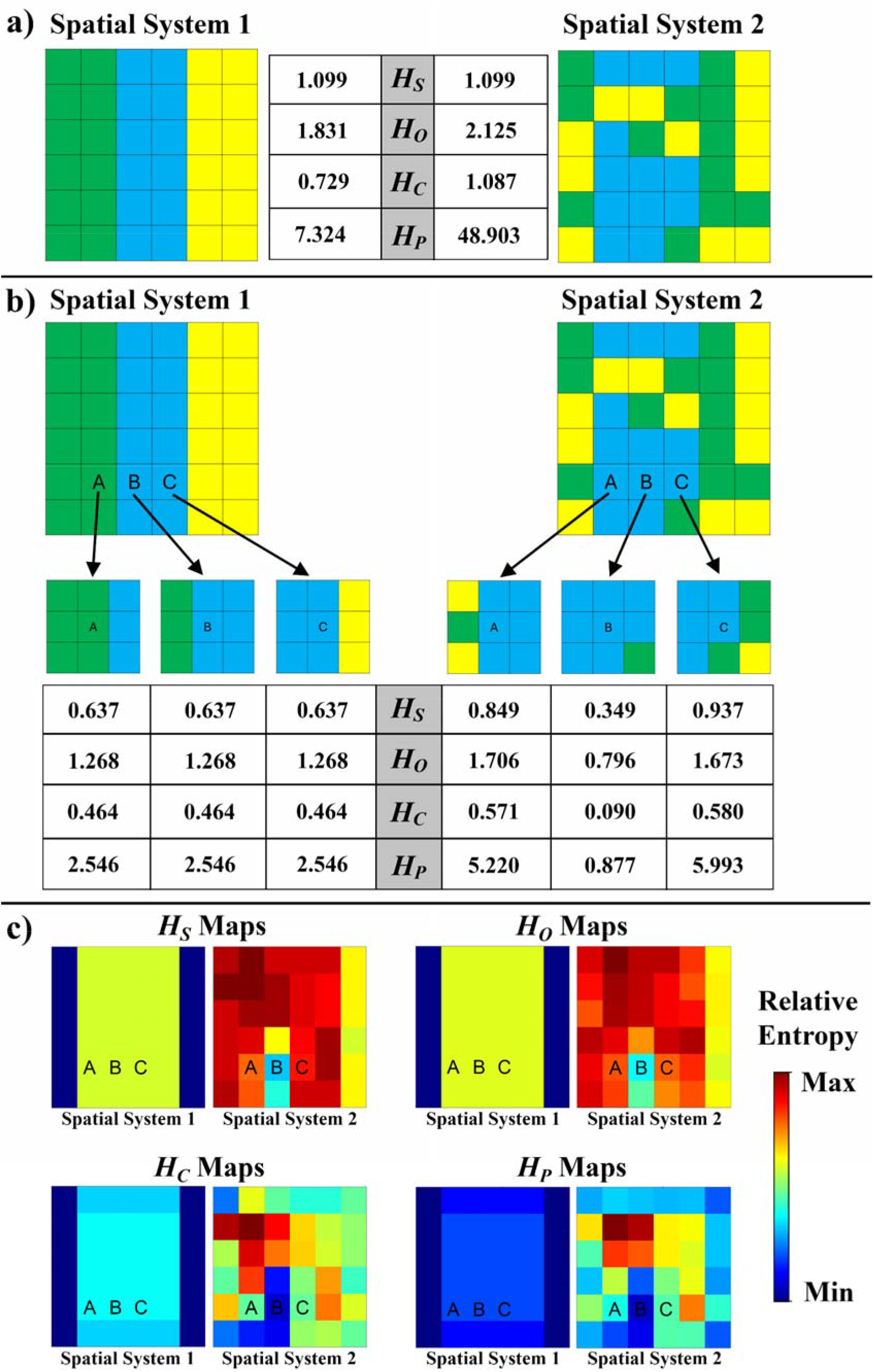
Spatial entropy is calculated for two spatial systems using two distinct approaches. In **a)**, the first approach is shown, where a single entropy value is calculated for each spatial

In Figure 1b, the same two spatial systems are divided into subareas and spatial entropy is calculated within each subarea. In this example, a subarea consists of a central element and its immediate neighbors, including diagonals (i.e., a 3×3 element grid). The entropy value calculated for the subareas centered on elements A, B, and C are shown for each entropy measure. While the location of elements A, B, and C are constant between the two spatial systems, the spatial patterns of the communities represented in subareas A, B, and C differ between systems. In spatial system 1 (left), all three subareas have highly similar spatial patterns (i.e., two columns of elements from one community and one column of elements from a different community). This is reflected in the entropy of the subareas; all three subareas have the same amount of spatial entropy, regardless of entropy measure. In spatial system 2 (right), the analogous subareas of elements A, B, and C have more disparity in spatial entropy. Across all four entropy measures, subareas A and C have relatively higher spatial entropy than the corresponding subareas in spatial system 1. The spatial entropy of subarea B is relatively lower in spatial system 2 compared to spatial system 1.

By assigning the entropy value of a subarea to the central element of the subarea, it is possible to create a map of the spatial entropy of a system. Figure 1c shows entropy maps yielded by the two spatial systems using each of the four entropy measures. By comparing the entropy maps of the two spatial systems, it is possible to identify specific subareas within the systems that have different entropy levels. Notably, *H*_*S*_ and *H*_*O*_ yield qualitatively similar entropy maps to one another, and *H*_*C*_ and *H*_*P*_ also resemble one another. All of the methods produce vertical columns of 0 entropy along the left and right borders of spatial system 1. This is because the subarea of the voxels in the most lateral columns of spatial system 1 feature voxels from only one community. Therefore, the subareas are completely homogenous, and diversity is minimum. Comparing regional entropy in spatial systems in this way is not possible when a single entropy value is calculated for the whole system, as in Figure 1a.

### Spatial Entropy in Simulated Community Mixing

Figure 2 depicts a simulation analysis of a 50×50 grid of elements that begins as highly ordered, with all elements of each community residing in the same quadrant. Spatial disorder is iteratively increased across 2.5 million iterations. In each iteration, an element is randomly chosen and its position is exchanged with a neighboring element (for more details, see Detailed Methods). In this way, the system gradually becomes less ordered as elements from each community diffuse across the entirety of the system. Whole-system entropy values, subarea-wise entropy maps, and the average entropy of all subareas in the grid were calculated in the system periodically throughout the diffusion process.

**Figure 2.**
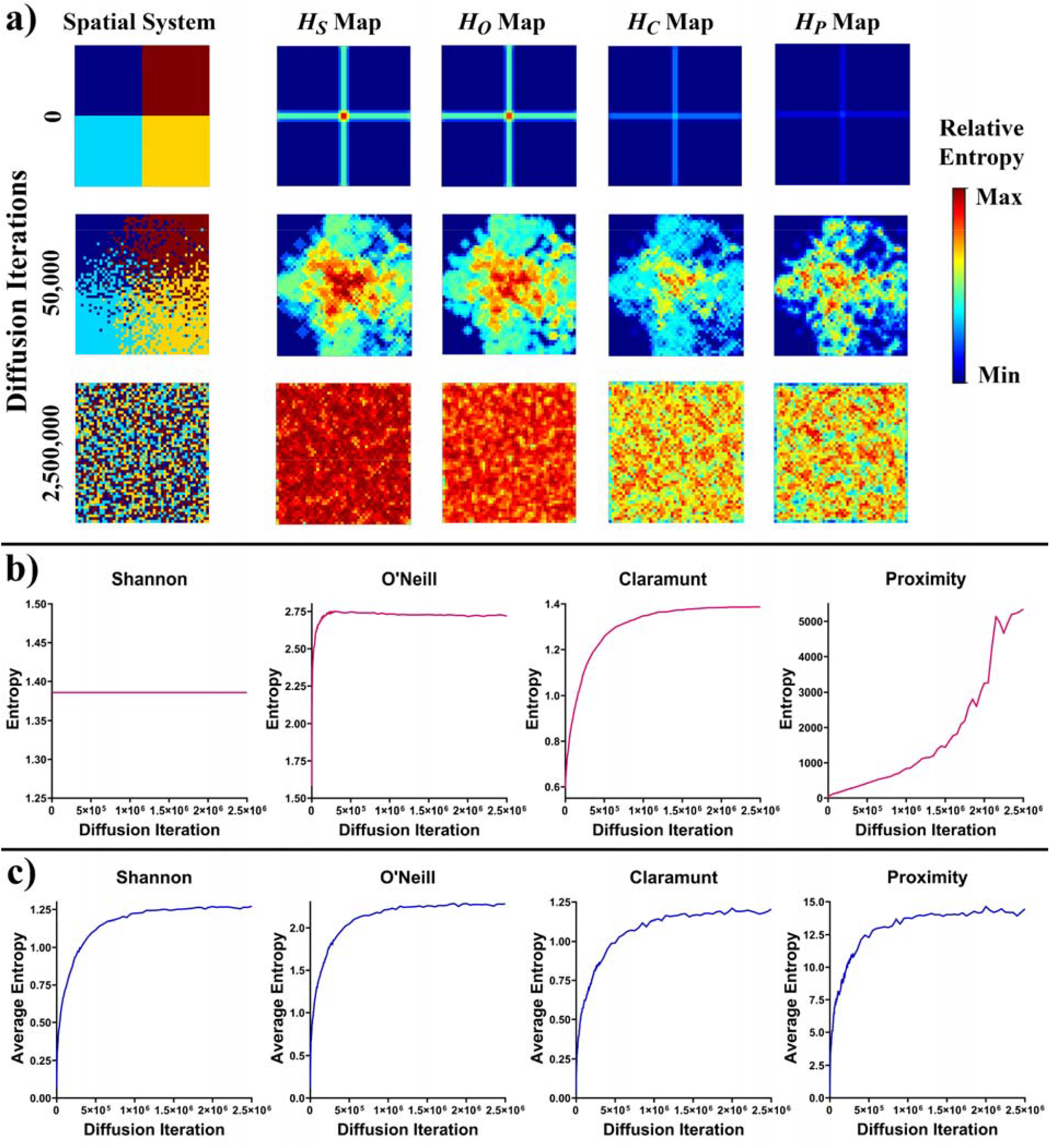
Results from simulated spatial patterns evolving from ordered to random. In **a)**, column 1 depicts a spatial system with four classes shown prior to any spatial mixing (row 1), after limited mixing (row 2), and after the system has been substantially mixed and approaches maximal spatial disorder (row 3). Columns 2-5 show the spatial entropy maps yielded when spatial entropy is calculated for each element subarea (all elements within two elements of the subarea center) according to four different spatial entropy measures. Color maps show relative entropy such that each entropy measure has a minimum of 0, but each measure has a unique maximum (maxima are 1.386, 2.738, 1.860, and 23.327 for *H*_*S*_, *H*_*O*_, *H*_*C*_, and *H*_*P*_, respectively). In **b)**, the entropy from calculating a single representative entropy value for the system is shown across iterations. In **c)**, the average entropy value across all elements in the system is shown across iterations. *H*_*S*_ – Shannon’s Entropy *H*_*O*_ – O’Neill’s Entropy *H*_*C*_ – Claramunt’s Entropy *H*_*P*_ – Proximity Entropy

In Figure 2a, three timepoints from the system are shown. The first timepoint is at baseline, prior to any spatial mixing. Spatial entropy is at a minimum regardless of entropy measure for this timepoint. The second timepoint (iteration 50,000) represents a period in the diffusion process during which entropy is quickly increasing. The final timepoint shows the state of the system after 2.5 million iterations. Figure 2b shows line plots corresponding to whole-system entropy throughout the full diffusion process. *H*_*S*_ never increases because the proportion of elements belonging to each category is constant throughout all iterations. *H*_*O*_ has a sharp increase in early iterations before flattening out. Both *H*_*C*_ and *H*_*P*_ show relatively gradual increases in whole-system entropy throughout the whole simulation with *H*_*P*_ not reaching a steady state until the end of the simulation. Figure 2c shows the average entropy across all subareas of the system. When measured this way, the differences between entropy measures are minimized with all approaching steady state near 1 million iterations. Supplementary Video 1 shows an animated version of this simulation wherein changes to the spatial pattern of the grid and corresponding changes to entropy maps and whole-system entropy values can be followed in detail.

### Applying Spatial Entropy to Functional Brain Network Data Structures

Functional brain networks were generated at the voxel level using blood oxygen level dependent (BOLD) signal measured with fMRI (for details, see Detailed Methods). Community detection algorithms (for details, see Detailed Methods) were used to identify communities of voxels that were more closely connected with each other than to voxels of other communities. Following community detection, each voxel had a community assignment. These community assignments were mapped into brain space according to the identity of each voxel. These three-dimensional community structure maps in brain space were the landscapes from which we generated entropy maps, with each brain scan yielding a distinct entropy map.

Figure 3 shows this process in two representative participants’ brain networks. In column one, the networks are shown as graphs in a non-Euclidean space. Nodes are colored according to the network community to which the node was assigned. Community assignments can be mapped into standard brain space allowing for assessment of community locations in the brain at the individual level. This is shown in column two, where each voxel from the graphs is represented in brain space. The colors of voxels in the community maps match the colors of the voxels in the graphs from column one. Because communities are unique to individual brains/networks, direct comparison across participants is not meaningful. Even in communities that bear a passing resemblance across participants – such as the turquoise voxels in the occipital lobe of both participant 1 and 2 – the communities have slight differences in voxel makeup along with differences in how the community is connected with the rest of the brain network.

**Figure 3.**
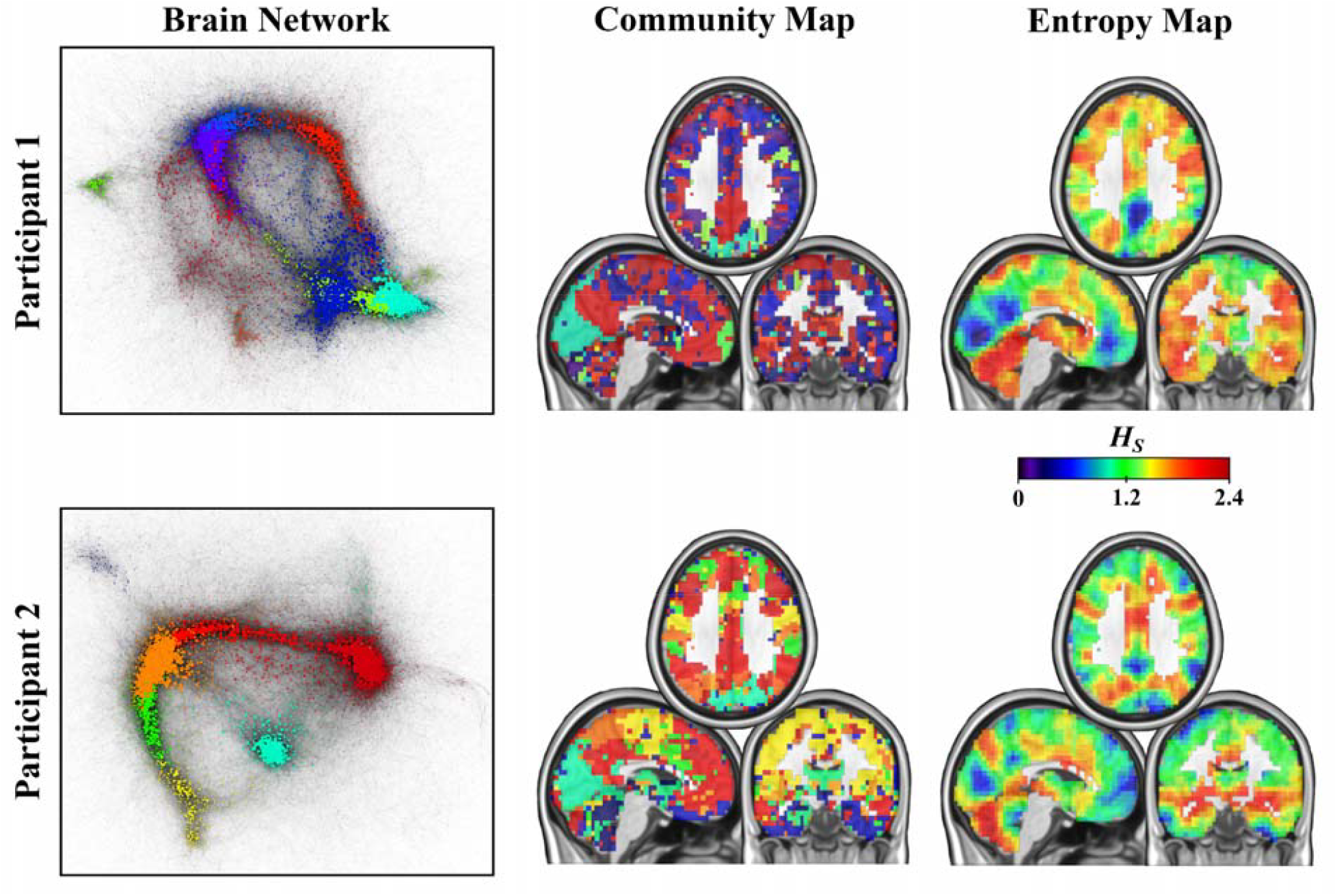
Two representative brain networks are shown. In column one, networks are shown in graph space, with nodes colored according to their community assignments. In column two, voxels are shown in brain space and are colored according to the same community assignments as column one. In column three, the *H*_*S*_ map corresponding to the community structure maps in column two are shown. *H*_*S*_ – Shannon’s Entropy

Therefore, by definition the communities are not the same and should not be compared as if they are. However, since community assignment data mapped to the brain is represented in a spatial structure, spatial entropy can be measured for each voxel according to the community assignments of voxels in the three-dimensional spatial subarea. Column three depicts the *H*_*S*_ maps that result from calculating spatial entropy for each voxel in the community maps from column two. In contrast to the community maps, entropy maps can be meaningfully compared across participants.

We note that it is possible and, in some cases, perhaps ideal to apply scaling parameters to entropy quantification. We explored the effects of several scaling parameters on entropy maps. These methodology of these supplemental analyses is presented in the Supplementary Material with results presented in Supplementary Figures 6-9. In general, these scaling approaches did not meaningfully change results compared to excluding the scaling parameters.

### Spatial Entropy Patterns Associated with Cognitive Processing

Entropy maps for resting-state and 2-back working memory scanning sessions were generated for n=22 healthy young adult subjects. Participant, task, and scanning details are provided in the Detailed Methods. Spatial entropy averaged across the whole brain was higher during 2-back scans relative to resting-state scans (Supplementary Figure 5). A primary goal for this work was to identify specific regions with differing entropy levels between rest and the working memory task. To account for higher global entropy in working memory scans compared to resting-state scans, entropy values were converted to Z-scores within scans for visualization.

Figure 4 shows entropy maps with each row corresponding to one of the four entropy measures. Column one shows group average Z-scored entropy maps from resting-state scans. Column two shows group average Z-scored entropy maps for 2-back working memory task scans. Column three maps the difference between columns one and two (i.e., rest entropy Z-score minus task entropy Z-score). Column four shows significant T-values for the rest versus task comparison using a paired t-test. Only regions that are part of significantly large clusters of voxels (minimum cluster size = 60 for *H*_*S*_, 71 for *H*_*O*_, 46 for *H*_*C*_, and 23 for *H*_*P*_). Results from *H*_*S*_, *H*_*O*_, and *H*_*C*_ were qualitatively very similar to each other in the first three columns while *H*_*P*_ produced entropy maps that were visibly very different. In particular, *H*_*P*_ maps tended to feature very prominent edge effects such that entropy was noticeably lower along the outer and inner edges of the brain parenchyma. The difference map for *H*_*P*_ was more similar to the other three measures, but differences between rest and task were weakest for *H*_*P*_.

**Figure 4.**
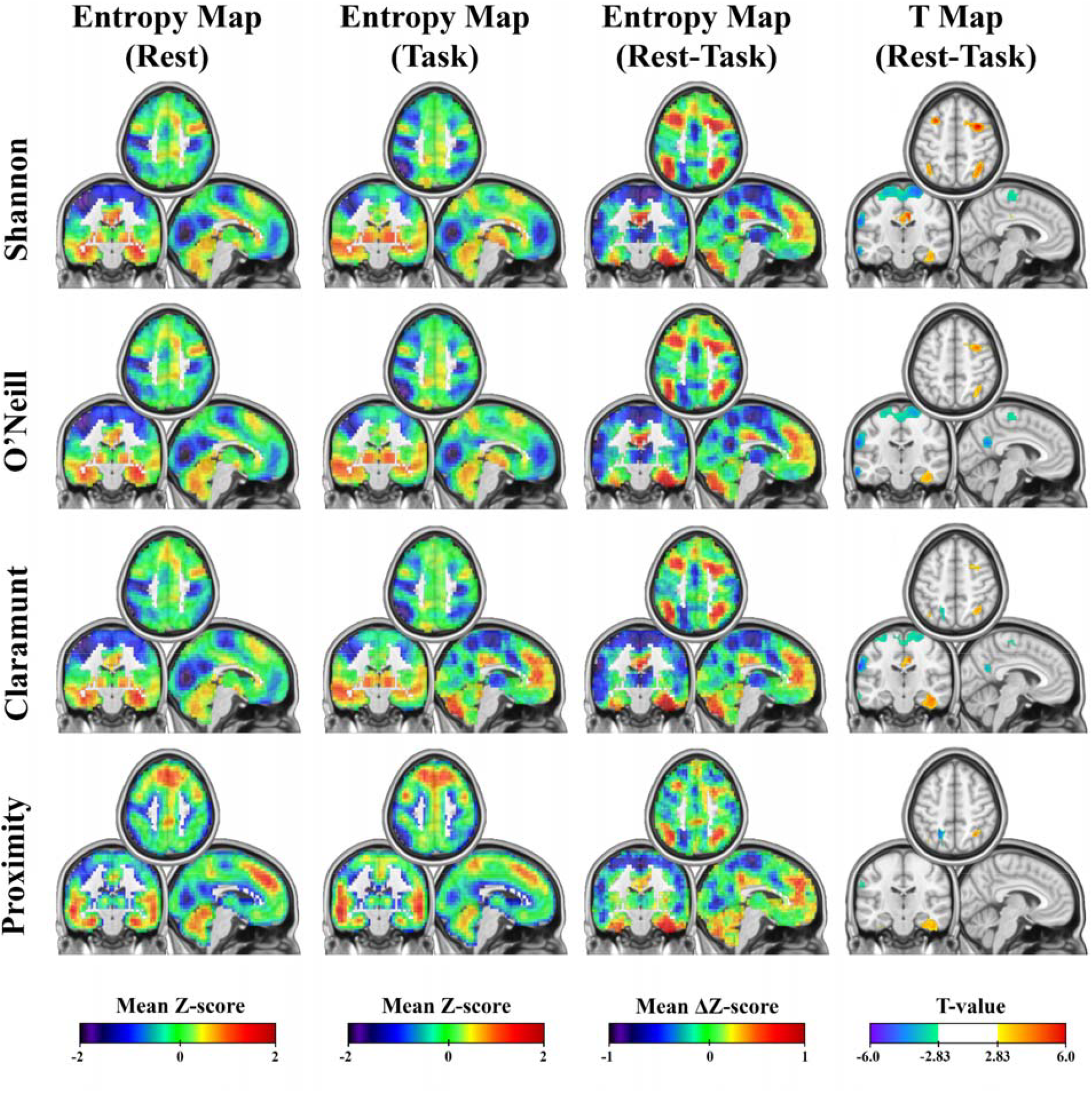
(Column 1) Sample average entropy maps for resting-state. Entropy values for each voxel subarea were converted to Z-scores within subject, then averaged across the sample. (Column 2) Results from the same sample average entropy mapping as in column 1, but for 2-back scans. (Column 3) The difference between entropy masks in columns 1 and 2. Note that color scale changes for this column. (Column 4) Maps of statistically significant t-values for paired comparisons of resting-state and 2-back. A height threshold of |T| > 2.83 (α = 0.005) with cluster correction for multiple comparisons (α = 0.05) was used to identify significant voxels. Positive T-values indicate higher entropy for rest compared to task. Negative T-values indicate lower entropy for rest than task. Each row corresponds to a different entropy measure. While the first three measures were highly similar, *H*_*P*_ was noticeably different from the other three measures. Shown slices have MNI coordinates X = -5, Y = -19, Z = 50

Supplementary Tables 1-4 give details of significant statistical results including cluster sizes and peak T-score locations. Across *H*_*S*_, *H*_*O*_, and *H*_*C*_, regions with higher entropy in rest compared to task included regions commonly associated with the CEN, although the statistical strength of this effect was strongest for *H*_*S*_. Other regions with higher entropy in rest compared to task were the midcingulate gyrus, left cerebellum, and right inferior temporal gyrus. Regions with lower relative entropy in rest compared to task included the posterior DMN, although the statistical strength of this effect was strongest for *H*_*O*_. Other regions with lower entropy during rest included the right inferior frontal gyrus, left and right supramarginal gyri, left and right medial pre-and postcentral gyri, left insula, and left middle occipital gyrus. *H*_*P*_ differences between rest and task were in similar locations and had similar effect directions as the other three measures but were weaker. Notably, when the *H*_*P*_ measure was used, rest and task entropy maps did not yield statistically significantly different clusters in the anterior CEN nor the posterior DMN.

### Working Memory Replication and Music Listening

Figure 5 compares results from the study above with an independent dataset of n = 5 individuals who completed several resting-state and 2-back scans within a single study visit (details in the Detailed Methods). Row one of Figure 5 is a reproduction of row one from Figure 4, and row two of Figure 5 shows results from the replication set. Results from the second working memory study were remarkably similar to results from the first study. In both studies, engagement of the CEN during working memory was associated with low entropy in this network. At rest, entropy in the precuneus was lower than during the task in both studies. This effect was statistically significant for *H*_*S*_ in the second study (a minimum of 91 voxels were required to survive cluster correction) despite not being statistically significant for *H*_*S*_ in the first study.

**Figure 5.**
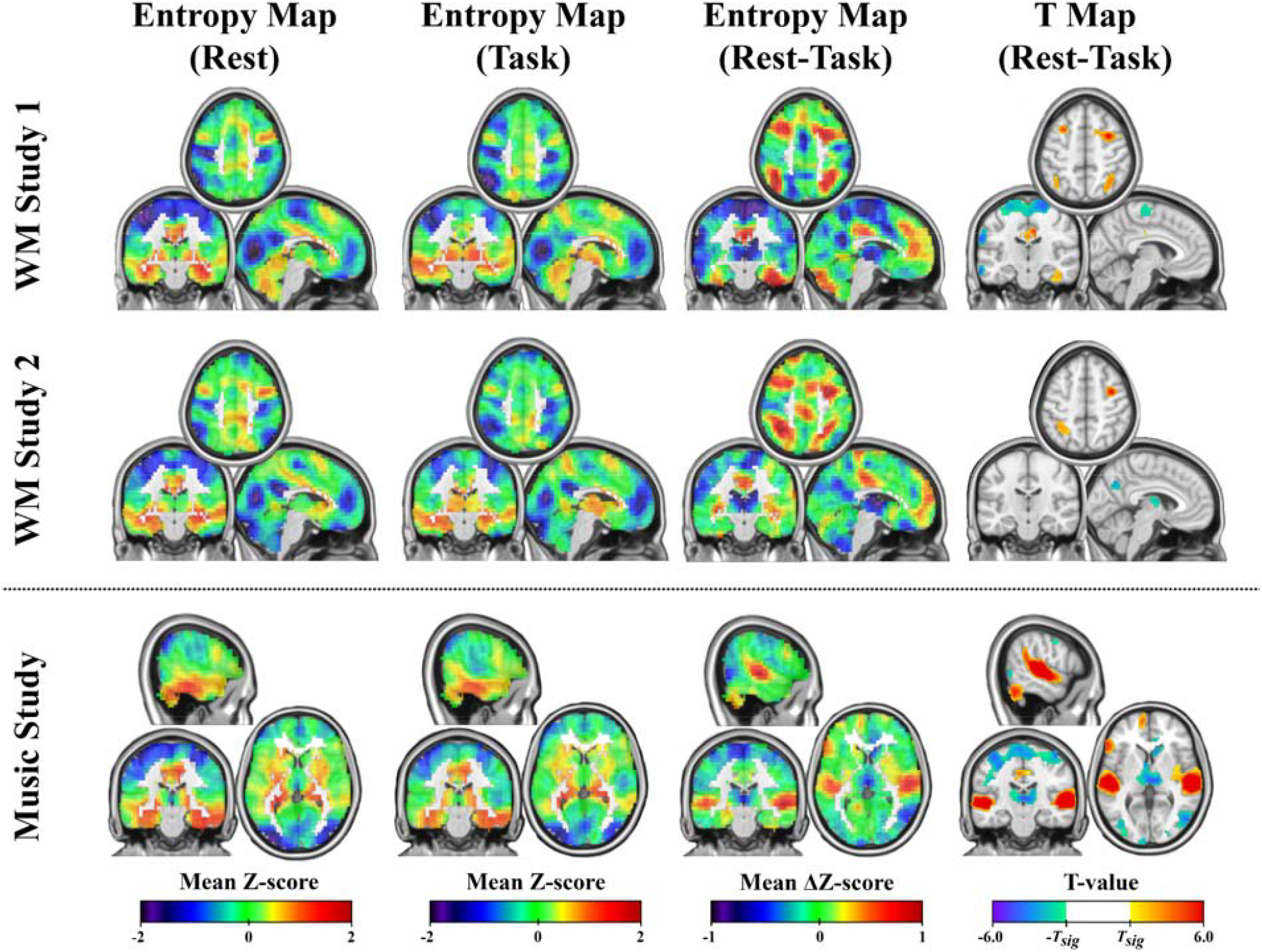
Sample average entropy maps for resting-state (Rest, column 1) and task (Task, column 2), the difference between rest and task (column 3), and maps of statistically significant t-values for paired comparisons of resting-state and 2-back (column 4). *H*_*S*_ was used for each row. Row **a)** shows results from the first working memory study, which were previously shown in Figure 4. Row **b)** shows results from the second working memory study. Results between the two studies were highly similar despite samples being independent, scanning protocols differing, and community detection algorithms differing. Row **c)** shows results from the music-listening tasks of study three. Of note, the auditory cortex had significantly lower entropy during the music listening tasks relative to resting-state scans. *T*_*sig*_ is variable by study and corresponds to the threshold for statistical significance for individual voxels. It is equal to approximately 2.83 in row one, 2.80 in row two, and 2.62 in row three. Slices from rows **a)** and **b)** have MNI coordinates X = -5, Y = -19, Z = 50 Slices from row **c)** have MNI coordinates X = 58, Y = -22, Z = 3 *WM – Working Memory*

Row three of Figure 5 demonstrates a completely separate application of *H*_*S*_. For this application, n = 20 healthy young adult participants underwent resting-state and music-listening scans (details in Detailed Methods). Resting-state entropy maps are shown in column one. Column two shows entropy maps for participants as they listen to songs of different genres in the MRI scanner. Column three shows the difference between resting-state and music-listening, and column four highlights significant differences between rest and task (a minimum of 62 voxels were required to survive cluster correction). The primary finding was that the left and right primary auditory cortices demonstrated lower spatial entropy during the music-listening task relative to resting-state.

## Discussion

The goal of this study was to introduce the use of spatial entropy to quantify the organization of brain network community structure. To achieve this goal, four spatial entropy methods were applied to simulated spatial systems and real functional brain network data. Using these spatial entropy metrics in three datasets, we demonstrated that engagement in a cognitive or perceptual task resulted in decreased entropy in expected brain regions. Importantly, we also observed changes in the local entropy of unexpected regions during tasks in each of the three studies. This highlights a key feature in spatial entropy analyses – they are entirely data-driven, can yield results that are consistent with known neurobiological principles, and may inspire new ideas about the functional (re)organization of the brain during different cognitive states.

Results from simulated spatial systems were included in this work to show simple examples of how spatial entropy can be quantified and interpreted at both the whole-system and subarea level. Figures 1 and 2 both highlight the inability of *H*_*S*_ to capture increasing spatial disorder at a whole-system level when the number of elements from each community is unchanged. This limitation was expected because *H*_*S*_ was not designed to account for the spatial distributions of system elements. This limitation motivated the development of the *H*_*O*_, *H*_*C*_, and *H*_*P*_ measures (O’Neill et al. 1988; Claramunt, 2005; Wang & Zhao, 2018), which were modified here to work with brain network data. Each of these methods successfully captured increases in system-level spatial entropy with the introduction of spatial disorder in our simulated data. When the focus of spatial entropy quantification shifted from the whole system to subareas, there were several occasions in which the four entropy measures disagreed about which subareas have higher or lower entropy due to the specifics of each method (Figure 1 and Supplementary Figure 2).

When applied to functional brain network data, three of the four entropy measures *(H*_*S*_, *H*_*O*_, and *H*_*C*_) had qualitatively similar entropy maps for both rest and task. *H*_*P*_ produced maps with an edge effect such that voxels in the center of large patches of gray matter had higher entropy than voxels at the edges of the brain or near the lateral ventricles, likely due to fewer opportunities for voxels in subareas to have touching borders. Across all measures, the CEN had lower relative entropy during 2-back scans compared to rest. In contrast, the posterior DMN had lower entropy during rest. While the difference maps (Figure 4) showed this effect across all entropy measures, only some measures had statistically significant differences. We interpret lower entropy during task as evidence of neural self-organization that is supporting cognitive demand. We also note that regions with entropy changes during task were not limited to the hypothesized DMN and CEN regions. At rest, lower entropy was seen in the supramarginal gyri, sensorimotor cortices, insula, and the occipital gyrus. While not part of the DMN, these regions have previously shown higher organization at rest to support sensory processing (Nelson et al., 2010; Oberhuber et al., 2016; Renier et al., 2010; Zhao et al., 2019). Importantly, findings were very closely reproduced in a second independent sample.

To show that spatial entropy is applicable beyond working memory, we applied *H*_*S*_ to a dataset (Wilkins et al., 2014) in which participants completed a resting-state scan and music-listening scans. As expected, entropy levels in the auditory cortices were lower when participants were listening to music, which is consistent with previous work showing that the auditory cortex is implicated in music-listening and sound-processing (Toader et al., 2023). We interpret the change in entropy as an increase in the spatial organization of the auditory cortices allowing for sound perception and processing. This finding provides evidence that spatial entropy is a valid measure of regional organization for a range of tasks and is not unique to any one task or dataset.

We note that we benefited from decades of task-activation-based work with fMRI and PET imaging (Belin et al., 1999; D’Esposito et al., 1995; Raichle et al., 2001), which led us to hypothesize that working memory engagement would reduce entropy in the CEN, and that listening to music would reduce entropy in the auditory cortices. While activation studies helped us to generate hypotheses, we note that this method is not just a surrogate measure of activation. It is a measure of the spatial organization of complex networks that addresses challenges to group-level community structure analysis. It is clear from Figures 4 and 5 that across all three studies, entropy differences between tasks were not limited to the regions that we knew to look for based on activation literature. We believe this highlights the potential for this method to not only confirm existing beliefs, but also to learn something new.

An important question following the results presented in this work is “which entropy measure is most appropriate for brain entropy maps?”. As shown in Supplementary Figure 2, *H*_*C*_ can fail when there is only one voxel from a given community in the subarea. For this reason, it may be best to avoid *H*_*C*_ for subarea-level spatial entropy analyses. We also discourage applying *H*_*P*_ in subareas due to its apparent bias towards having lower entropy around the edges of the brain as shown in Figure 4 (also see edge effects in the simulated system in Figure 2a). We suggest applying either *H*_*S*_ or *H*_*O*_ for spatial entropy analyses at the subarea level. In theory, *H*_*O*_ is superior because it accounts for the spatial arrangement of voxels within subareas while *H*_*S*_ does not. In practice, the two measures yielded highly similar results in this work. We believe that calculating entropy for each voxel based on its subarea (i.e., within a relatively small space) helps to overcome the theoretical shortcomings of *H*_*S*_ for spatial systems.

This study acknowledges several limitations. The first is the relatively small sample size of the datasets in this study (n = 22, 5, and 20). Additionally, as with any brain network analysis, we had to make several choices in network modeling, such as the specific approach to quantifying edge strength, the process of binarizing edges, and which community detection algorithm to use. An exhaustive comparison of how all of these factors influence entropy maps is beyond the scope of this work. In the future, it would be prudent to further explore these factors in focused studies on ideal data collection and modeling parameters for spatial entropy analyses. It very well may be that there is no “correct” approach and that the most appropriate combination of parameters is dependent on the research question. We also note that while we did not observe any impact of scaling parameters on our entropy maps, it is possible that these factors may have a more meaningful influence on other datasets.

To our knowledge, this is the first study to investigate the organization of functional brain networks by identifying community structure and calculating spatial entropy at the voxel level. This approach provides a fully data-driven approach to analyzing brain network community structure at the group level. We believe that this method will prove successful in studying a wide variety of cognitive processes as well as brain disorders, including but not limited to cognitive decline in older adults and emotional response in psychiatric disease. We also note that we have not shown that high entropy is inherently “bad” or “good”. While results in this work suggest that low entropy supports cognitive processing, it is possible that in other contexts low entropy in key regions may contribute to inflexibility in brain networks, leading to excessive rumination and depressive symptoms. Finally, while this work focused on fMRI-based functional brain networks, there should be no reason why the same approach taken in this work cannot be applied to analyses of other network representations of spatial systems.

## Detailed Methods

### Datasets and Study Samples

This work includes data from three independent functional neuroimaging studies. Supplementary Figure 1 provides an overview of the three studies including the participant samples and tasks completed during scanning for each study.

The primary dataset used in this work is from a study that is described in detail in previous work (Mayhugh et al., 2016; Moussa et al., 2014). From the original study, the present work utilized each participant’s structural MRI scan along with a resting-state and a 2-back functional MRI scan from healthy 24-35 year old participants (n=22). All participants were recruited via local advertisements or by word-of-mouth and were compensated for participation. The Wake Forest University School of Medicine Institutional Review Board approved the study. All participants gave written informed consent. For more details about recruitment and inclusion/exclusion criteria for the first working memory study, see the Supplementary Material.

The second dataset used in this work has also appeared in prior work (Rzucidlo et al., 2013). In this study, n=5 healthy participants aged 23.6 ± 1.67 years underwent structural MRI scanning along with five alternating resting-state and 2-back functional scans (for a total of 10 scan runs) in a single session. This data was used in the present study to determine whether findings from the first working memory study were replicable in an independent sample with a different scanning protocol. The Wake Forest University School of Medicine Institutional Review Board approved the study. All participants gave written informed consent.

The third dataset used in this work has also been described in detail previously (Wilkins et al., 2014). This study included n=20 healthy participants aged 20-31 years old. Participants underwent structural MRI scans along with resting-state and music-listening functional scans. The music study was included in the present work to assess whether spatial entropy measures are capable of identifying functionally meaningful spatial patterns in community structure data aside from working memory engagement (i.e., listening to an auditory stimulus). This study was approved by The Wake Forest University School of Medicine Institutional Review Board. All participants provided written informed consent.

### Image Acquisition

For the first working memory study, image acquisition included a T1-weighted structural scan, a resting-state functional scan, and a 2-back functional scan. For the resting-state scan, participants were instructed to fixate on a cross projected onto a screen. For the 2-back scan, participants viewed a series of letters presented one at a time. When a new letter was presented, participants were instructed to indicate whether or not the currently-presented letter was the same as the letter that had been shown two letters prior. The task lasted 6 minutes and had 120 trials.

Each trial lasted for 3,000 ms, with a letter being presented for the first 300 ms followed by a blank screen for 2,700 ms. Participants could respond at any point during the trial using a button box. Participants practiced the task for one minute outside the scanner and an additional minute inside the scanner before image collection began. The original study included older adults that were excluded from analyses in the present work because they had low performance on the 2-back task (Mayhugh et al., 2016). MRI data were obtained on a 3T Siemens Skyra equipped with a 32-channel head coil, a rear projection screen, and both left and right hand response boxes. High-resolution (0.98 x 0.98 x 1.0 mm) T1-weighted structural scans were acquired in the sagittal plane using a single-shot 3D MPRAGE GRAPPA2 sequence (TR = 2.3 seconds, TE = 2.99 ms, flip angle = 9°). BOLD-weighted image sequences were acquired in the transverse plane using an echo-planar imaging sequence (resolution = 3.75 x 3.75 x 5.0 mm, acquisition time = 6 minutes and 20 seconds, TR = 2.0 seconds, TE = 25 ms, flip angle = 75°, 35 slices per volume, 187 volumes).

For the second working memory study, image acquisition included a T1-weighted structural scan, five resting-state functional scans, and five 2-back functional scans. Resting-state and 2-back scans were collected in alternating order. The resting-state and 2-back protocol for this study were the same as the first working memory study. All scans for this study were collected on the same MRI scanner as the first working memory study. A foam padded birdcage head coil was used to minimize movement artifacts. High-resolution (1.2 x 1.2 x 1.2 mm) T1-weighted structural scans were acquired in the sagittal plane using a single-shot 3D MPRAGE GRAPPA2 sequence (TR = 1.9 seconds, TE = 2.95 ms, flip angle = 9°). BOLD-weighted functional images (resting-state and 2-back) were collected using an echo-planar imaging sequence (resolution = 3.75 x 3.75 x 5.0 mm, acquisition time = 4 minutes and 20 seconds, TR = 2.0 seconds, TE = 25 ms).

For the music study, image acquisition included a T1-weighted structural scan, a resting-state functional scan, and music-listening functional scans. Participants listened to several genres of music in several distinct music-listening scans. Participants had their eyes closed for both the resting-state and the music-listening scan runs. To ensure that each music genre was represented by the same scan length, all song recordings were edited to be five minutes long. Scans for this study were collected with a 1.5T GE twin-speed LX scanner with a birdcage 12-channel head coil. High-resolution (1.2 x 1.2 x 1.2 mm) T1-weighted structural scans were acquired in the sagittal plane using a single-shot 3D BRAVO sequence (TR = 11.936 seconds, TE = 5 ms, flip angle = 12°). BOLD-weighted images were collected using a single-shot, gradient-recalled, echo-planar imaging sequence (resolution = 3.75 x 3.75 x 5.0 mm, acquisition time = 5 minutes, TR = 2.0 seconds, TE = 40 ms).

### Image Preprocessing

Given that the data from the three studies used here all are from different time periods over the past two decades, different preprocessing steps were used for each study in accordance with the best practices at the time the data for these studies were collected. We considered rerunning preprocessing steps for all scans; however, the objective of this study is to provide a framework for evaluating the entropy of community structure landscapes, not to evaluate how various preprocessing steps affect brain network community structure. Therefore, we have used the original preprocessing steps (described below) and feel that replication of findings between the two working memory studies serves to imply robustness of our technique to preprocessing specifics. A separate parametric study evaluating combination of different preprocessing steps is beyond the scope of this initial study describing spatial entropy methodology, though it would be a prudent future direction.

For the first working memory study, initial image preprocessing steps were completed using SPM8 (http://www.fil.ion.ucl.ac.uk/spm/). FSL fMRIB (Smith et al., 2004) was used to preprocess the second working memory and music studies. For all studies, structural images were skull-stripped, segmented into grey matter, white matter, and cerebrospinal fluid maps, and normalized to Montreal Neurological Institute (MNI) space. The first 6-10 volumes of functional scans were discarded to allow signal to achieve equilibrium. Functional images were slice time corrected and realigned to the first volume. The previously defined warp to MNI space was then applied to functional images. A band-pass filter (0.009-0.08 Hz) was applied to remove physiological noise and low-frequency drift. To remove global signal, the mean signals for whole brain, white matter, and CSF were regressed out of functional time series along with six degrees of freedom movement parameters. For the first working memory study, the motion scrubbing procedure had been introduced (Power et al., 2014), so this procedure was used to remove volumes with motion artifacts coupled with BOLD signal change.

### Network Generation

Functional brain networks were generated for each scan. Each network node represented a voxel. Each network edge represented a functional connection between voxels. Functional connections were quantified according to the Pearson correlation of the BOLD signal between each pair of voxels.

For the first working memory study, edge weights were thresholded and binarized following an algorithm that resulted in networks in which every node was included in the giant component. Network density was equal across subjects following the equation *S* = log(*N*) / log(*k*), where *N* is the number of nodes, *k* is average degree, and *S* = 2.5 as empirically determined by prior work (Hayasaka & Laurienti, 2010) and because this value has been used in prior functional network analyses of this data (Mayhugh et al., 2016). To avoid generating fragmented networks, a variation of the Minimum Spanning Tree algorithm was implemented.

First, a network with *S* = 2.5 using standard thresholding (maintaining only the strongest positive correlation values) was generated. The network was divided into its components. At this point, it would be possible to connect all components with the strongest shared edge between the largest connected component and every other component in turn; however, this would arbitrarily increase the density of networks. Therefore, in an iterative fashion, the density threshold was raised (forcing generation of sparser networks) by 1/(2^15^), and again the network was divided into its components. Each component was connected to the largest connected component through the strongest positive between-component connections. With this process, the number of edges in the thresholded network decreases faster than the number of disconnected components increases. This process was continued until the number of edges needed to include all nodes in one connected component did not result in exceeding the *k* needed to satisfy *S* = 2.5, resulting in connected networks with the same density across all scans.

For the second working memory and music studies, edges were thresholded and binarized such that the most strongly correlated edges were included in the binarized network while satisfying density *S* = 2.5. We did not require that all nodes were contained in a single connected component.

### Network Community Detection

Functional brain networks tend to yield distinguishable modules or “communities” of nodes that are more connected with each other than they are with other nodes in the network. This property is quantifiable by the modularity measure *Q* (Newman & Girvan, 2004). For the first working memory study, *Q* was optimized using a dynamic Markov process called “stability” (Delvenne et al., 2010). All parameters were kept at default values to find the community (i.e., module) partitions that maximized *Q*. Because this optimization algorithm has a stochastic element, the algorithm was run 10,000 times. The partition with the highest *Q* out of the 10,000 iterations was used as the representative community mapping for the brain network. This process was carried out for each scan such that every individual had a unique community mapping for resting state and for 2-back. For the second working memory and music studies, community assignments were optimized using the QCut algorithm (Ruan & Zhang, 2008).

### Entropy Methods – Original Shannon and Batty

In the original work introducing entropy *(H)* as a measure of uncertainty (also called diversity or information) in a discrete system (Shannon, 1948), *H* is calculated as

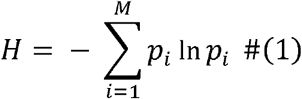

where *M* is the number of possible states and *p*_*i*_ is the probability of a system being in state *i*. From this equation it follows that if each state is equally likely, *H* is maximized for the system. If probabilities of observing different states are not equal, *H* is lower as there is less uncertainty about the state of the system as one or more states have higher probability of occurring.

Conceptually, entropy has many potential applications, including quantifying the extent to which the spatial organization of a system is structured or disordered. However, using entropy in a spatial context requires a modified approach. This is because as originally proposed, *H* is solely concerned with the probability distribution of variables - not the spatial distribution of variables. To address this limitation, a spatial form of entropy was introduced (Batty, 1974). To quantify spatial entropy, Batty proposed partitioning a large area into smaller subareas. The probability of observing categorical variables within subareas could then be quantified according to an equation that is roughly equivalent to Equation 1. From this point, Batty’s use of spatial entropy (which was originally proposed to study large scale ecological biomes) ceases to be applicable to brain network analysis. For brain network analysis, the key contribution from Batty was the intuition to divide large spatial systems into smaller subareas. This innovation allows entropy to be quantified at a local level, which makes it possible to begin considering the spatial structure of large systems.

Batty’s approach of partitioning a space into subareas serves as the starting point for the method proposed in this paper. The full 3-dimensional volume of each participant’s community map was partitioned into many smaller subareas of voxels. Subareas were partitioned according to spatial proximity within the full brain volume. Spatial entropy was then quantified within each subarea for each participant according to the entropy methods described below.

### Entropy Methods – Spatial Shannon

The first measure of spatial entropy for brain network data that was evaluated in this work was based on Shannon’s *H*. Essentially, this measure quantifies the Shannon entropy for each subarea. For each voxel, neighboring voxels within a certain distance (see *Subarea Size*) of the reference voxel were included in entropy calculation. That is, a subarea was centered around each voxel. Because this measure focuses on voxel subareas rather than the full brain, this approach allows for distinguishing between brain regions of relatively high or low spatial entropy. Therefore, it is distinct from the original *H* proposed by Shannon and is instead a spatial form of Shannon’s entropy (*H*_*S*_). Following this implementation, spatial Shannon’s entropy *H*_*S*_ is calculated for each voxel as

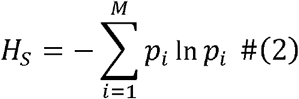

where *M* is the total number of communities in the network and *p*_*i*_ is the probability of observing a voxel belonging to community *i* in the voxel subarea. *H*_*S*_ is highest if each voxel represents a unique community (i.e., no community has more than one voxel representative). *H*_*S*_ is lowest if a single community dominates the voxel subarea.

### Entropy Methods – O’Neill

Another approach to quantifying spatial entropy was introduced by O’Neill (O’Neill et al., 1988) and will be referred to as O’Neill’s entropy (*H*_*O*_). Like *H*_*S*_, *H*_*O*_ begins by dividing an area into subareas. The methods differ in their calculations of entropy within subareas. Whereas Shannon’s entropy is concerned with the proportion of categories represented in subareas, O’Neill’s entropy is concerned with pairings of categories that share borders within the subarea. *H*_*O*_ is calculated for each voxel subarea as

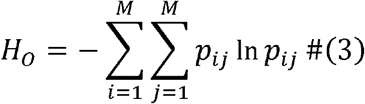

where *p*_*ij*_ is the probability of observing a contiguous pairing of voxels from community *i* and community *j*. Therefore, *H*_*O*_ is highest when each combination of contiguous pairings of communities are equally represented. *H*_*O*_ is lowest when only one community is present in the subarea. If multiple communities are present in the subarea, *H*_*O*_ is lowest if all contiguous edge pairings are the same (e.g., a checkerboard pattern would have minimal *H*_*O*_ for a subarea with two communities present).

### Entropy Methods – Claramunt

Another quantification of spatial entropy was introduced by Claramunt (Claramunt, 2005) and will be referred to as Claramunt’s entropy (*H*_*C*_). As with the previous entropy measures, we begin our implementation of *H*_*C*_ for each voxel by identifying a subarea of voxels around the center voxel. The unique aspect of *H*_*C*_ is that it incorporates the distance between elements of the same versus different categories within the subarea. *H*_*C*_ is very similar to *H*_*S*_ but adds in a fractional scaling factor that adjusts the entropy contribution of each community according to the distance between elements of the same versus different communities. That is, *H*_*C*_ is calculated for each voxel as

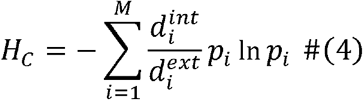

where 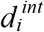 is the average distance between voxels within community *i* and 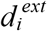 is the average distance between voxels belonging to community *i* and voxels belonging to other communities. *H*_*C*_ is maximized when communities are evenly spatially distributed across the subarea. *H*_*C*_ is minimized when the distance between voxels of different communities is much larger than the distance between voxels of the same community (i.e., when voxels of the same community are spatially clustered in the subarea).

### Entropy Methods – Proximity

A final measure of spatial entropy is provided by Wang and Zhao (Wang & Zhao, 2018) and is referred to as Proximity entropy (*H*_*P*_). Conceptually, *H*_*P*_ is a combination of *H*_*O*_ and *H*_*C*_ as it adds a scaling factor to *H*_*S*_ that accounts for shared edges as well as distance. *H*_*P*_ is calculated as

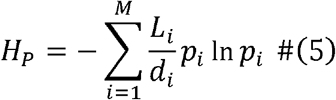

where *L*_*i*_ is the total number of touching edges between voxels in community *i* and voxels of other communities and *d*_*i*_ is the sum of the average distances between the centers of different communities. To calculate *d*_*i*_, the center of each community must be calculated by finding the average position of voxels belonging to the community in the subarea. The average distance between voxels of community *i* and the center of every other community is calculated. The sum of these average distances (one average distance per non-*i* community) is *d*_*i*_. In theory, *d*_*i*_ can be less than one and substantially inflate the entropy of a subarea. To avoid outlying subareas complicating analyses, *d*_*i*_ is set to 1 for all subareas with *d*_*i*_ < 1. *H*_*P*_ is high when many voxels of different communities are touching and the centers of communities in the subarea are close to each other. *H*_*P*_ is low when voxels mostly touch other voxels of the same community and the centers of communities are spatially distant in the subarea.

### Simulated Spatial Systems

The primary motivation for this work was to demonstrate how different entropy measures translate to brain network analysis and to assess their biological relevance. However, as this is the first work to apply these methods to brain network community structure analysis, we believe it is important to clearly demonstrate how the different entropy measures differ in theory so that future investigators can make informed decisions about which measure to apply in their analyses. Therefore, we generated several examples of simulated 2-dimensional spatial systems to demonstrate how spatial entropy differs depending on the measure used.

First, we demonstrate how entropy measures differ when calculating a single entropy value for a large spatial system. This approach is representative of how spatial entropy measures are typically used in ecology. We then demonstrate how entropy can be calculated for each distinct subarea within the larger spatial system, resulting in entropy maps for the system. This approach is representative of our proposed application of spatial entropy measures to brain network analysis.

We then demonstrate how the four entropy measures have theoretical differences in terms of what constitutes a highly entropic spatial pattern using several examples of simple 2-dimensional subareas. The spatial subareas in these examples are smaller and generally simpler than the 3-dimensonal subareas in brain network analysis – the intent of these examples is to impart on the reader intuition as to how the relative entropy of different spatial patterns can differ depending on the entropy measure that is applied.

Finally, we provide an example of a spatial system that begins as highly ordered but gradually becomes more entropic based on the following rules. The system begins in a 50×50 grid of elements, with each element belonging to one of four categories. An equal number of elements belongs to each category, and elements of the same category are arranged in quadrants at the beginning of the simulation. The system then gradually changes in an iterative fashion such that randomly selected elements trade positions with their neighbors. The initially obvious boundaries between quadrants of categories begin to fade until eventually the system appears maximally disordered. We calculate the entropy of the system at each iteration according to the ecology-like approach to studying spatial entropy (i.e., calculating one value for the system). We also generate entropy maps with subareas centered on each element in the grid and calculate the whole-system entropy by averaging the entropy values of the elements.

### Subarea Size

An important, modifiable parameter for all entropy measures is the size of the subarea surrounding each voxel. We generated entropy maps for three different subarea sizes for each entropy measure. For the three subarea sizes, voxels were included in the subarea if the Euclidean distance to the subarea center voxel was within one, two, or three voxels. For subarea size one, the maximum number of voxels in subareas was 7 (the center voxel and six immediate neighbors located at a Euclidean distance of one voxel). For subarea size two, there were up to 33 voxels in subareas. For subarea size three, there were up to 123 voxels in subareas. While it is possible to calculate entropy on subareas extending four or more voxels from the subarea center, such subareas would provide little spatial specificity. Therefore, subareas of this size were not assessed in this work. Based on the entropy maps generated for these different subarea sizes, we focused on subareas extending three voxels from the center voxel as these results appeared smoother and most interpretable while maintaining a degree of spatial specificity.

### Comparing Entropy Maps Across Tasks

Resting-state and task entropy maps were compared in the three studies in this work using paired t-tests in SPM12. Due to observed differences in average subarea entropy, grand mean scaling was use for each entropy map in each study prior to statistical testing.

For the first working memory study, there was a single rest scan and a single task scan per participant. Therefore, each participant had exactly one pair of scans in the model.

For the second working memory study, each participant had five rest scans and five task scans. The first rest scan was compared to the first task scan, the second rest scan was compared to second task scan, and so on for all five scans for each participant. Participant identity was included as a covariate to account for repeated measures.

For the music scans, each participant had one rest scan and six task scans. The same rest scan was included in the model six times, with each image being paired with one of the six task scans for the participant. Again, participant identity was included as a covariate to account for repeated measures.

Differences in entropy values between rest and task were considered statistically significant for voxels with *p* < 0.005. To correct for multiple comparisons, only significant clusters of voxels were retained. The number of voxels required for a cluster to be significant was unique to each study and is listed in the associated results section.

## Data and Code Availability

Code used for generating entropy maps is openly available at https://github.com/rlyday/brain-network-spatial-entropy.

