## Supplement for "Spatial entropy of brain network landscapes: a novel method to assess spatial disorder in brain networks"

*Sample Recruitment and Inclusion/Exclusion Criteria*

Exclusion criteria for the parent study included history of alcohol or drug abuse/dependence, self-reported use of illicit drugs in the past 3 months, or recent/ongoing (within the last 6 months) Axis I disorders according to DSM IV definitions. Participants were also excluded if they reported more than one binge drinking (≥5 drinks for men and ≥4 drinks for women in less than 2 hours) occasion in the past 3 months, if they failed to pass a breathalyzer (i.e., if the participant had > 0.00 BAC based on measurements from an Intoxilyzer S—D5 breath alcohol screen, [www.alcoholtest.com](http://www.alcoholtest.com/)) at the beginning of any study visit, or if they had a positive test for cocaine, THC, opiates, amphetamines, or benzodiazepines (measured by an Alere iCassette 6 –panel drug screen, www.alere.com) at the beginning of any visit.

Participants were also excluded if they had Modified Mini Mental State Exam score equal to or below 80, if they had active neurological dysfunction that might impact cognitive ability (e.g., schizophrenia, Alzheimer’s disease, attention deficit-hyperactivity disorder, Parkinson’s disease, epilepsy, or prior history of stroke), or if they used antipsychotic or antiepileptic medications. Participants were also excluded if they had previous serious central nervous system trauma (i.e., sub- or epidural hematomas), previous brain surgery, or loss of consciousness greater than 5 minutes. Antidepressant use was allowed if they had been in use for over 2 months prior to study entry and the participant did not have active depression as measured by the Center for Epidemiological Studies Depression Scale (CES-D) or Profile of Moods Survey (POMS). Individuals with Body Mass Index (BMI) < 18 or > 35 were also excluded, as were individuals with insulin-dependent type I diabetes, high blood pressure without at least 1 year of stable treatment, those with positive pregnancy tests, left-handedness, corrected visual acuity < 20/40, or impaired hearing.

The second working memory study (i.e., the replication dataset) was originally designed to examine the stability of brain network topology within and between resting state and working memory task states. Participants were healthy adults recruited from around Forsyth County, North Carolina. Participants that had any implants, devices, or objects that would interfere with the fMRI procedure were excluded from the study.

The music listening study was designed to investigate the effects of listening to music of different genres on functional brain network connectivity. Participants were recruited based on their preferred musical genre such that each genre had similar numbers of participants. All of the participants were right-handed, English speaking, color-sighted, had normal hearing, and did not have any neurological disorders.

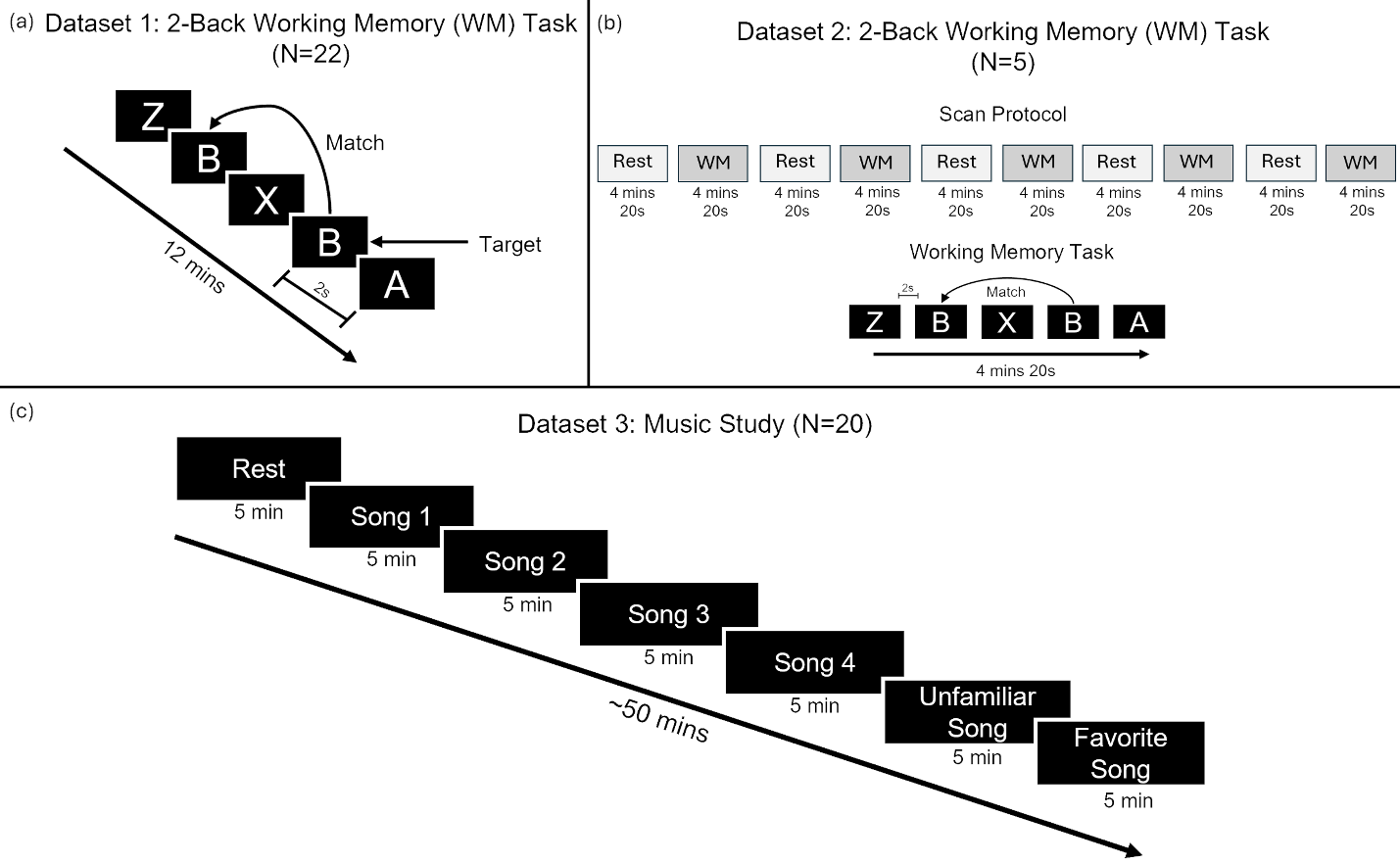

**Supplementary Figure 1.** Overview of datasets used for this study. (a) 2-back working memory task completed in 22 participants over 6 minutes 20 seconds. (b) The N-back working memory task alternated with resting scans with a brief delay between each scan in 5 participants. Rest and 2-back scans were completed five times each. (c) 20 participants listened to six songs, each presented randomly based on genre preference from a pre-study questionnaire. These four songs of different genres and the unfamiliar song were consistent across participants, whereas the favorite song was participant-specific.

*Entropy in Simulated Subareas*

Supplementary Figure 2 depicts four pairs of 3x3 element subareas along with the corresponding subarea entropy calculated according to the four spatial entropy measures. Each pair of subareas features a slight difference in the subarea composition. These slight changes often result in changes in entropy, but in several examples the direction of change from one subarea to its partner differs depending on the entropy measure.

In Supplementary Figure 2a, swapping the positions of the center yellow element and the blue element to its left results in an increase in entropy according to *H_O_*, a decrease in entropy according to *H_C_* and *H_P_*, and no change in entropy for *H_S_*. *H_O_* is lower on the left subarea because there is only one combination of touching elements present, whereas the right subarea has two combinations. *H_C_* and *H_P_* are both higher on the left side because the two communities are maximally interspersed, resulting in low distances between elements of different communities and the centroids of different communities. *H_S_* is unchanged because the number of elements in each community is equal between the two subareas.

Similarly, in Supplementary Figure 2b, swapping the positions of the green element and the center blue voxel result in an increase in the subarea entropy according to *H_C_* and *H_P_*, a decrease in entropy for *H_O_*, and no change in entropy for *H_S_*. Again, *H_O_* deviates from *H_C_* and *H_P_* because it is solely concerned with touching pairs of elements, and there are more combinations of touching pairs present in the subarea on the left. On the right, the single green element is more interspersed with the large body of blue elements, which raises *H_C_* and *H_P_*. The proportion of voxels in each community is equal between the left and right subarea, so *H_S_* is also equal between the left and right subarea.

In Supplementary Figure 2c, the subarea includes 9 different communities. For a subarea with 9 elements, this should result in maximal entropy. On the left, replacing the navy element on the top right of the subarea with a yellow element decreases entropy for *H_S_*, *H_O_*, and *H_P_*, all as expected. However, this particular situation reveals something important about *H_C_*. For the spatial system on the left, *H_C_* is equal to 0. Recall from Equation 5 that the scaling factor introduced by Claramunt features in its numerator the average distance between elements of the same class. In the spatial system on the left, each element is the only element of its class in the subarea. Because there are no other elements of the same class, the distance is equal to 0, and the entropy values of class *m* is scaled to 0. This example reveals that classes with only one element in a subarea cannot contribute directly to the subarea entropy (i.e., the entropy contribution to *H_C_* from a community with only one element in the subarea is equal to 0).

In the final example, Supplementary Figure 2d shows the same left spatial pattern as Figure Supplementary Figure 2c. However, in the right spatial pattern, the yellow element replaces the pink element on the upper left of the subarea rather than the navy element on the upper right (as in Supplementary Figure 2c). *H_S_* is higher on the left subarea because the number of elements is equal across communities on the left, but not on the right. *H_O_* is the same on the left and right, because every touching pair of elements have only a single shared border. As in Supplementary Figure 2c, *H_C_* is higher on the right subarea because it has a community (yellow) with two elements in it whereas the subarea on the left has a single element from each community. *H_P_* is higher on the right because the centroid of the yellow community is drawn more towards the center of the subarea in the subarea on the right. This reduces *d_i_*, the sum of distances between different communities, which is the denominator in the scaling factor in the *H_P_* calculation. Smaller total distance between communities therefore increases *H_P_*.

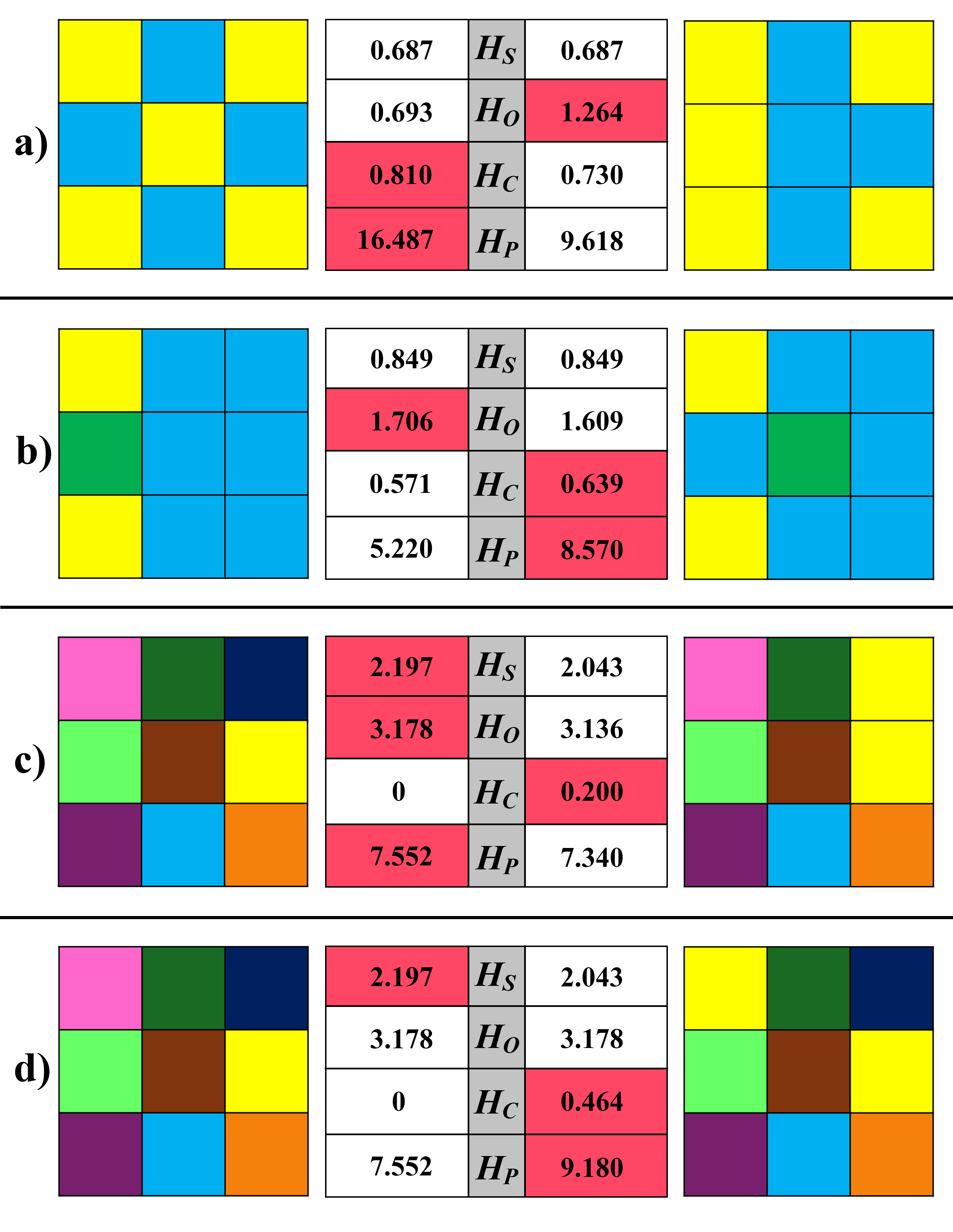

**Supplementary Figure 2.** Four pairs of spatial patterns are shown in **a-d**. The entropy of each pattern is quantified according to four different entropy measures. For each measure, the pattern with the higher entropy is highlighted in red. In **a)**, *H_S_* is equal between the two patterns, *H_O_* is higher on the right pattern, and *H_C_* and *H_P_* are higher on the left. In **b)**, *H_S_* is again even between the two patterns and the higher entropy pattern differs in H_O_ compared to *H_C_* and *H_P_*. In **c)**, *H_S_*, *H_O_*, and *H_P_* are higher for the left pattern, but *H_C_* is equal to 0 for the left pattern and is higher in the right pattern. In **d)**, *H_S_* is higher on the left while *H_C_* and *H_P_* are higher on the right. *H_O_* is equal between the two patterns.

*H_S_* – Shannon’s Entropy

*H_O_* – O’Neill’s Entropy

*H_C_* – Claramunt’s Entropy

*H_P_* – Proximity Entropy

*Supplementary Video 1*

An animated representation of the spatial mixing simulation (from Figure 2 in the main text) is available at the following link: <https://www.youtube.com/watch?v=YuZddQpXp_E>

*Impact of Subarea Size in Brain Network Data*

The majority of brain network analyses in this work focus on the first Working Memory study. In this study, larger subarea sizes had a smoothing effect that resulted in larger patches of voxels with relatively high or low entropy in entropy maps. This effect is demonstrated in Supplementary Figure 3 using *H_S_* as the entropy measure for resting-state scans. Supplementary Figure 4 shows this effect for the other three entropy measures. The smoothing effect facilitated interpretation of entropy maps and subsequent analyses comparing tasks for all four entropy measures. Therefore, subareas extending three voxels (i.e., the largest subarea size we assessed) are the focus of the remainder of this work.

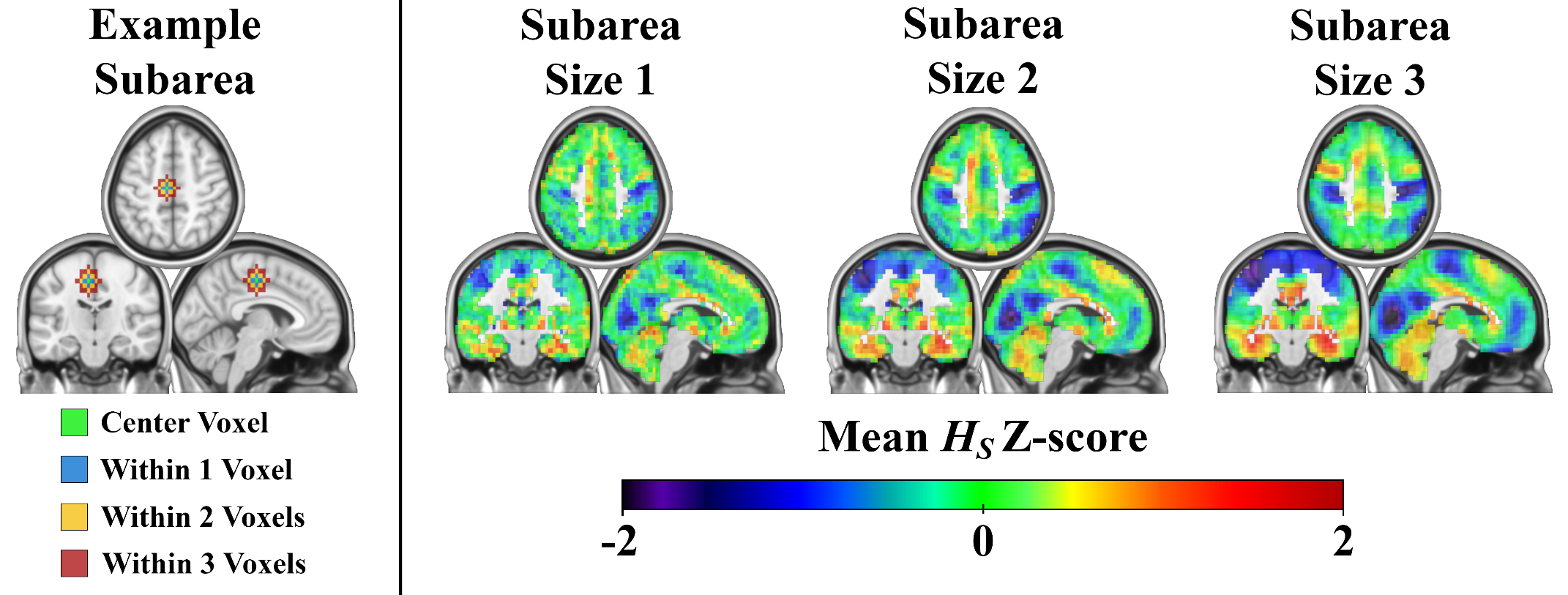

**Supplementary Figure 3.** Spatial Shannon entropy maps calculated on subareas of different sizes. Column 1 shows an example voxel subarea in three planes with voxels colored to indicate the size of the subarea required for voxels to be included in the entropy calculation for the center (green) voxel. Columns 2-4 show mean Z-scored *H_S_* maps for subareas extending one, two, and three voxels from the center voxel, respectively. Entropy values for each voxel subarea were converted to Z-scores within subject, then averaged across the sample for each subarea size. Increasing subarea size had a smoothing effect on entropy maps that made differences between regions of relatively high and low spatial entropy more visibly apparent.

Slices shown here have MNI coordinates X = -5, Y = -19, Z = 50

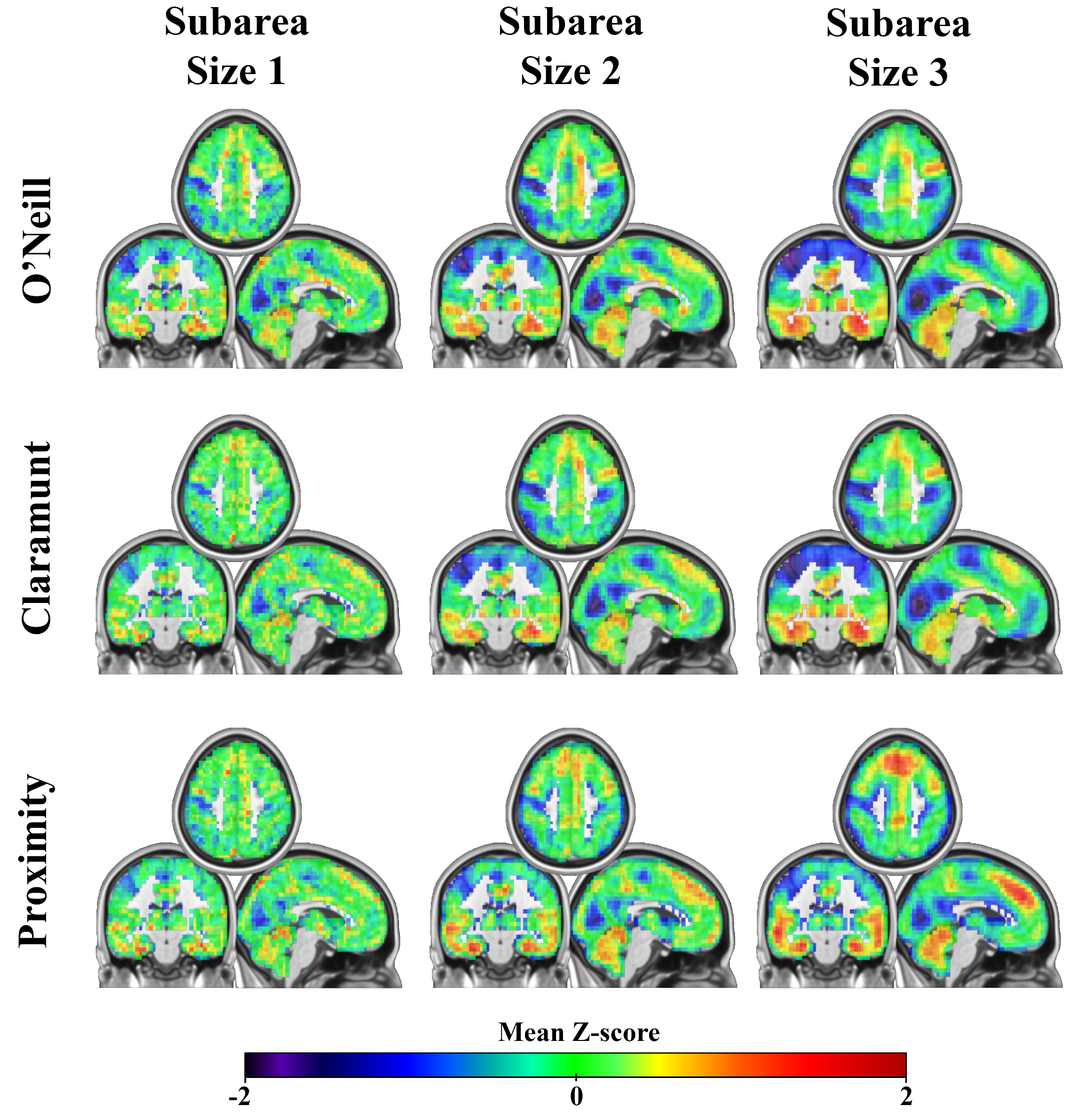

**Supplementary Figure 4.** Spatial entropy maps calculated on subareas of different sizes for the O’Neill, Claramunt, and Proximity entropy measures. From left to right, subareas included the center voxel and voxels within one, two, and three voxels of the center voxel, respectively. Entropy values for each voxel subarea were converted to Z-scores within subject, then averaged across the sample for each subarea size. Increasing subarea size had a smoothing effect on entropy maps that made differences between regions of relatively high and low spatial entropy more visibly apparent.

Shown slices have MNI coordinates X = -5 , Y = -19, Z = 50

*Measuring Whole System Entropy*

Paired t-tests assessed statistical significance of the difference when the entropy of the whole brain was calculated for resting-state and 2-back for each entropy measure. With this approach, *H_S_* (*p* = 0.188), *H_O_* (*p* = 0.359), and *H_C_* (*p* = 0.200) did not significantly differ between rest and task. *H_P_* was significantly higher during the 2-back task compared to rest (*p* < 0.001). These results are depicted in Supplementary Figure 5a.

We also compared whole brain entropy in a second approach that involved first calculating entropy for each voxel according to its neighborhood, then finding the average entropy value across all voxels in the brain. With this approach, all four entropy measures indicated that entropy was higher during the 2-back task compared to rest, all with *p* < 0.001 in paired t-tests. These results are depicted in Supplementary Figure 5b.

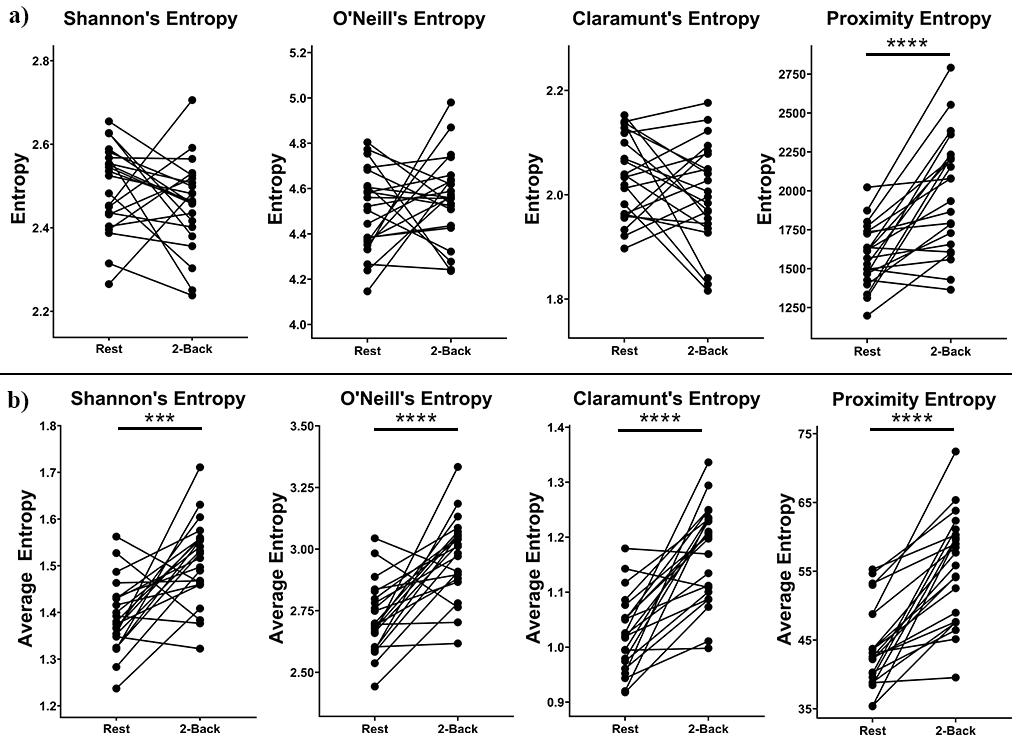

**Supplementary Figure 5.** Plots in **a)** show the whole-system entropy for resting-state versus 2-back scans according to each measure. Each pair of points represents a participant with lines connecting resting-state and 2-back scans. Of the four entropy measures, only Proximity entropy was significantly different between resting-state and 2-back scans. Plots in **b)** show the average subarea entropy values across the whole brain for each participant. Here, 2-back scans had higher entropy than resting-state scans for each of the four entropy measures.

*** indicates *p* < 0.001

**** indicates *p* < 0.0001

*Entropy Scaling by Distance*

Distance between voxels are intrinsic aspects of *H_C_* and *H_P_*. However, distance is not an explicit factor in *H_S_* or *H_O_*. Therefore, the effect of scaling entropy to account for distance between voxels in these two measures was assessed. In addition to generating entropy maps according to Equations 3 and 4, entropy maps were generated according to a scaling rule that made voxels nearer to the center of the subarea have more weight than voxels at the exterior of the subarea.

In the first implementation of *H_S_* (Equation 3 of the main paper), *p_i_* is calculated by counting the number of voxels belonging to each community *i* and dividing that number by the total number of voxels in the subarea. As an equation, *p_i_* is calculated as

$$\begin{aligned} p_{i}=\frac{v_{i}}{v_{n}} \#\left( S1 \right) \end{aligned}$$

where *v_i_* is the number of voxels in community *i* and *v_n_* is the number of voxels in the subarea. In this implementation, each voxel in the subarea has an equal weight.

In the distance-scaled implementation, each voxel is weighted according to the distance to the center voxel of the subarea. Voxel weights were equal to 1/*d*, where *d* is the Euclidean distance (in voxels) from the center voxel. The center voxel and its immediate neighbors had weight 1. A voxel located three voxels away from the center of a subarea would have *d*=3 and would be weighted as 1/3. Therefore, in the distance-scaled implementation, *p_i_* is calculated by summing the weights of voxels in community *i* and dividing that number by the sum of all voxel weights in the subarea. This is calculated as

$$\begin{aligned} p_{i}^{\left( D \right)}=\frac{\sum_{x=1}^{v_{i}} \frac{1}{d_{x}}}{\sum_{x=1}^{v_{n}} \frac{1}{d_{x}}} \#\left( S2 \right) \end{aligned}$$

where (*D*) denotes that *p_i_* is scaled by distance to the center of the subarea, *v_i_* is the number of voxels in the subarea belonging to community *i*, *v_n_* is the total number of voxels in the subarea, and *d_x_* is the distance from voxel *x* to the center of the subarea. The distance-scaled implementation of Shannon’s entropy (*H_S_^(D)^*) is calculated as

$$\begin{aligned} H_{S}^{\left( D \right)}=-\sum_{i=1}^{M} p_{i}^{\left( D \right)}\ln p_{i}^{\left( D \right)} \#\left( S3 \right) \end{aligned}$$

The distance-scaled implementation of O’Neill’s entropy (*H_O_^(D)^*) is similar but accounts for the average distance of pairs of voxels rather than individual voxels. That is, the contribution of each pair of nodes *ij* is weighted according to the average distance of node *i* and node *j* from the center of the subarea. As with *H_S_^(D)^*, the center voxel is assigned distance 1 along with its immediate neighbors.

*Entropy Scaling by Community Size*

The influence of the size of communities (i.e., the number of voxels in each community) was also assessed for each entropy measure. The reasoning for this was that some communities are more likely than others to have many voxels in a subarea because the number of total voxels in each community is uneven. There are two conflicting interpretations for how to account for this characteristic of brain networks.

One interpretation is that if a voxel belonging to a small community is observed in a subarea, it would be more surprising than if a voxel from a large community was observed in the subarea. Therefore, voxels belonging to smaller communities should have more weight than voxels from larger communities. A subarea with relatively high numbers of voxels from small communities should have higher entropy than a subarea with mostly voxels belonging to large communities.

A second interpretation relates to practical issues that arise with small communities. If communities are smaller than the size of the subarea (i.e., there are more voxels in subarea *n* than there are voxels in community *i*), the theoretical minimum entropy for subareas that include voxels of community *i* is increased. In other words, because it is impossible for subareas with voxels from small communities to have only a single community present, it is impossible for these subareas to have 0 entropy. This is true even if all voxels of community *i* are spatially clustered within the subarea (i.e., even if community *i* is as spatially ordered as possible given its small number of voxels). Therefore, even when the spatial structure of small communities may be highly ordered, the subarea entropy will tend to be high if community size is not accounted for. Given this practical issue, this interpretation would suggest that larger communities should contribute more to the entropy of the subarea than smaller communities because smaller communities tend to inappropriately raise the minimum entropy of subareas.

The most reasonable interpretation and scaling implementation may depend on the specific research question and the data available. Therefore, we suggest an adjustable parameter, τ, that modifies the influence of community sizes such that smaller communities or larger communities can be more influential in subarea entropy calculations depending on the specified τ. The community-size-scaled implementation of Shannon’s entropy (*H_S_^(MS)^*) is calculated as

$$\begin{aligned} H_{S}^{\left( MS \right)}=-\sum_{i=1}^{M} \left| \tau-p_{i}^{b} \right|\times p_{i}^{n}\ln p_{i}^{n} \#\left( S4 \right) \end{aligned}$$

where *p_i_^b^* is the proportion of voxels in the whole brain belonging to community *i*, *p_i_^n^* is the proportion of voxels in the subarea belonging to community *i*, and τ is the community size tuning parameter. When τ is equal to 1, the entropy is scaled such that the contributions of smaller communities to the entropy of the subarea is weighted more than the contributions of larger communities. When τ is equal to 0, the opposite is true, and the contributions of larger communities are weighted more than the contributions of smaller communities. Therefore, τ = 1 is in line with the first proposed interpretation of community size (i.e., that it is surprising to observe voxels from a small community in the subarea). τ = 0 is in line with the second proposed interpretation (i.e., that the uncertainty of larger communities may be of more practical interest for applications of spatial entropy to brain network analysis). To determine how this scaling factor impacts findings, we generated entropy maps for both τ = 1 and τ = 0.

*Entropy Scaling by Number of Communities*

Different brain networks often have different numbers of communities. As the number of communities in a network increases, the theoretical maximum entropy for each subarea also increases as long as the number of voxels in subareas is greater than or equal to the number of communities in the network. Subareas of different sizes have different theoretical maximums in entropy because the number of communities that can possibly be represented is uneven across subareas. Therefore, it may be appropriate to scale subarea entropy values by either the number of communities in a network or by the number of voxels in the subarea – whichever value is lower. For example, this “normalized” entropy value can be computed for Shannon’s entropy (*H_S_^Norm^*) as

$$\begin{aligned} H_{S}^{(Norm)}=\frac{H_{S}}{\ln\left( \min\left( M_{b},v_{n} \right) \right)} \#\left( S5 \right) \end{aligned}$$

where *H_S_* is the originally calculated spatial Shannon’s entropy according to Equation 3, *M_b_* is the number of communities in the brain network, and *v_n_* is the number of voxels in the subarea. While *H_S_* is shown in Equation S5, the same normalization can be applied to the other three entropy measures.

*Entropy Scaling by Subarea Size*

Subareas across the brain include non-uniform numbers of voxels. This is because subareas centered in the outermost regions of the brain or near the lateral ventricles have neighboring voxels that do not include brain tissue. In the original implementation of *H_B_* proposed by Batty, a factor *T_g_* scales entropy for each category *i* according to the size of the subarea. To assess how this parameter modifies entropy maps, entropy maps were generated for each entropy measure. Just as in Batty’s original implementation, *T_n_* was used to divide *p_i_* (*p_ij_* for *H_O_*) within the log functions where *T_n_* is the number of voxels in the subarea.

*Impact of Scaling Parameters in Brain Network Data - Results*

Supplementary Figure 6 demonstrates the effect that the different scaling parameters had on *H_S_* rest and task entropy maps, difference maps, and T-value maps. Analogous maps for the other three entropy measures are shown in Supplementary Figures 7-9. In general, the influence of added scaling parameters did not qualitatively change results compared to the original entropy measures without the added scaling.

Scaling entropy contributions of voxels in the subarea according to their distance from the center of the subarea did not have a noticeable impact on *H_S_* or *H_O_*. This scaling parameter was not applied to *H_C_* or *H_P_* because distance is already incorporated into their calculations.

Scaling by module size with the first approach (τ = 1) did not noticeably change any mappings for any of the entropy measures.

Scaling by module size with the second approach (τ = 0) produced the most different mappings relative to mappings without additional scaling. For this scaling approach, entropy was noticeably higher in the dorsomedial prefrontal cortex, particularly during resting-state scans. This difference was evident in difference maps and T-value maps. This scaling approach was also the only one without significant differences between tasks in the right fusiform gyrus, left mid temporal gyrus, and lateral sensorimotor cortex.

Scaling by the number of modules in the brain network seemed to slightly reduce differences in entropy between task and rest, but it did not change spatial patterns for any entropy measures.

Scaling by subarea size resulted in edge effects that are visible in the first two columns of Supplementary Figures 6-9. The entropy in subareas centered along the exterior of the brain and near the lateral ventricles were relatively low due to having fewer voxels. These edge effects were consistent between rest and task for *H_S_* and *H_O_*, so they did not noticeably alter the difference maps or T-value maps. For *H_C_* and *H_P_*, scaling by subarea size substantially altered difference maps and T-value maps.

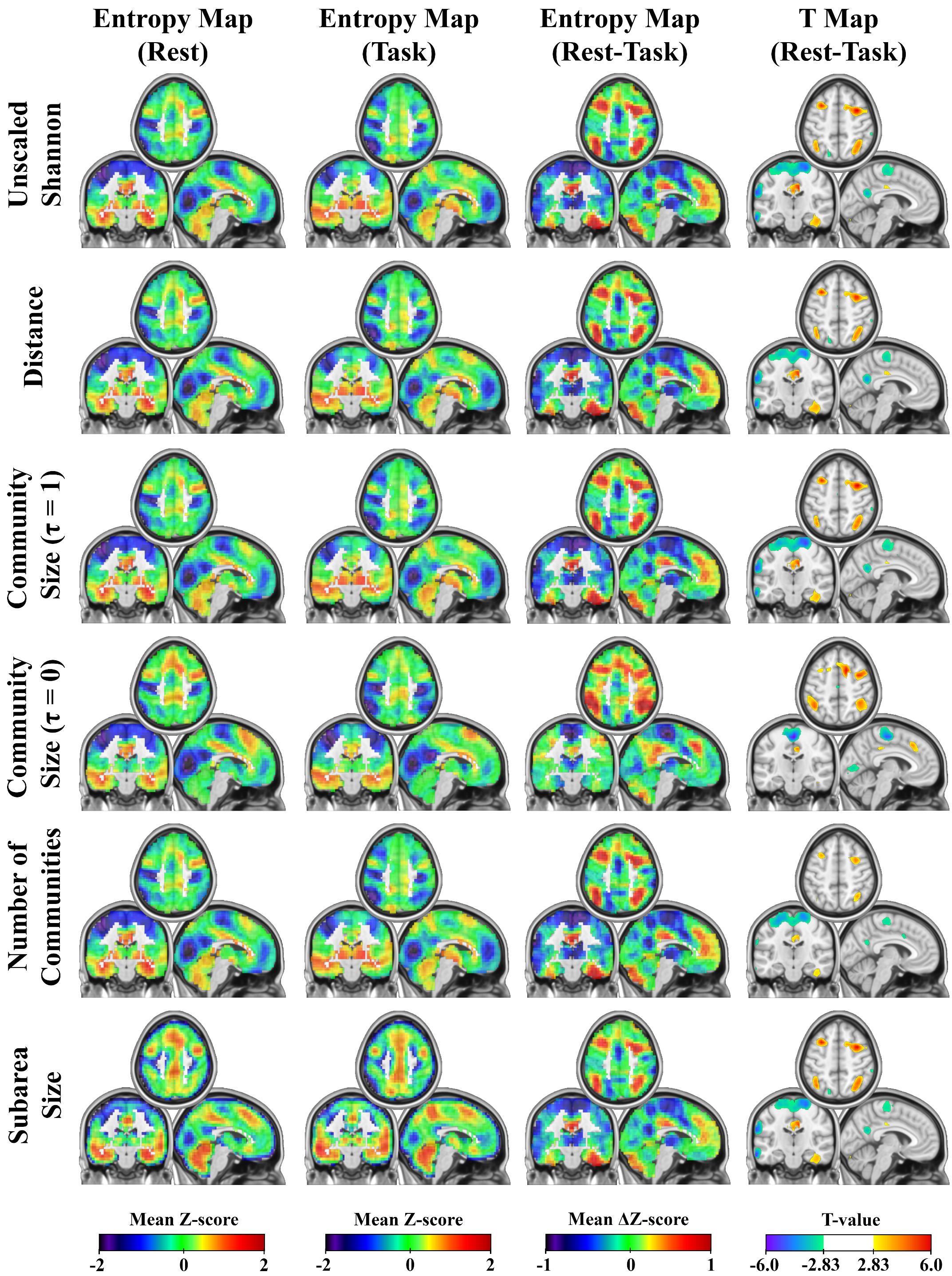

**Supplementary Figure 6.** Sample average entropy maps for resting-state (Rest, column 1) and 2-back (Task, column 2), the difference between tasks (column 3), and maps of statistically significant (without multiple comparisons correction) t-values for paired comparisons of resting-state and 2-back (column 4). Note that color scales change between the entropy maps (columns 1 and 2) and the difference maps (column 3). Each row corresponds to a different set of scaling parameters in comparison to Shannon’s spatial entropy without scaling (row 1).

Shown slices have MNI coordinates X = -5, Y = -19, Z = 50

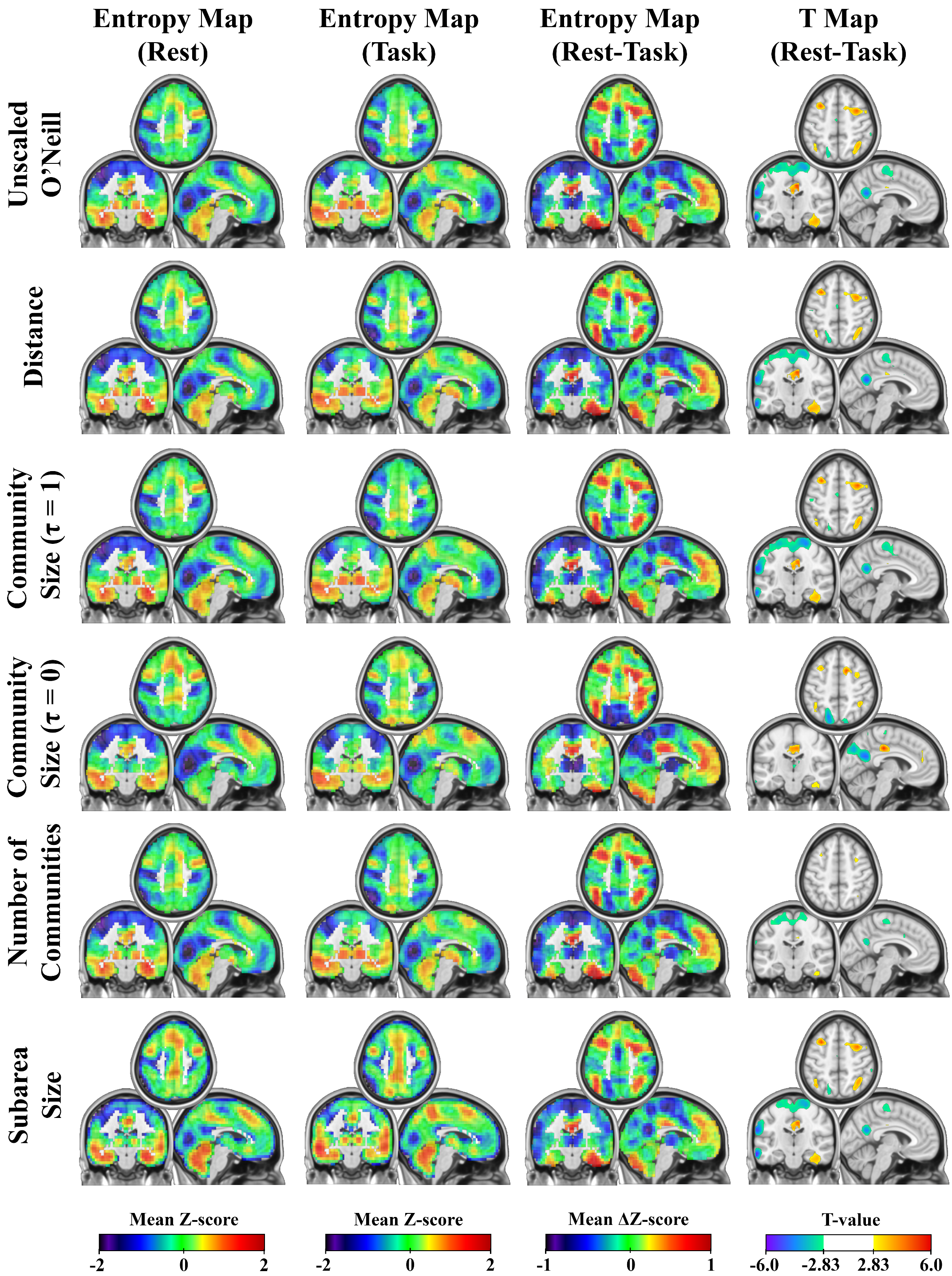

**Supplementary Figure 7.** Sample average entropy maps for resting-state (Rest, column 1) and 2-back (Task, column 2), the difference between tasks (column 3), and maps of statistically significant (without multiple comparisons correction) t-values for paired comparisons of resting-state and 2-back (column 4). Note that color scales change between the entropy maps (columns 1 and 2) and the difference maps (column 3). Each row corresponds to a different set of scaling parameters in comparison to O’Neill’s entropy measure without scaling (row 1).

Shown slices have MNI coordinates X = -5 , Y = -19, Z = 50

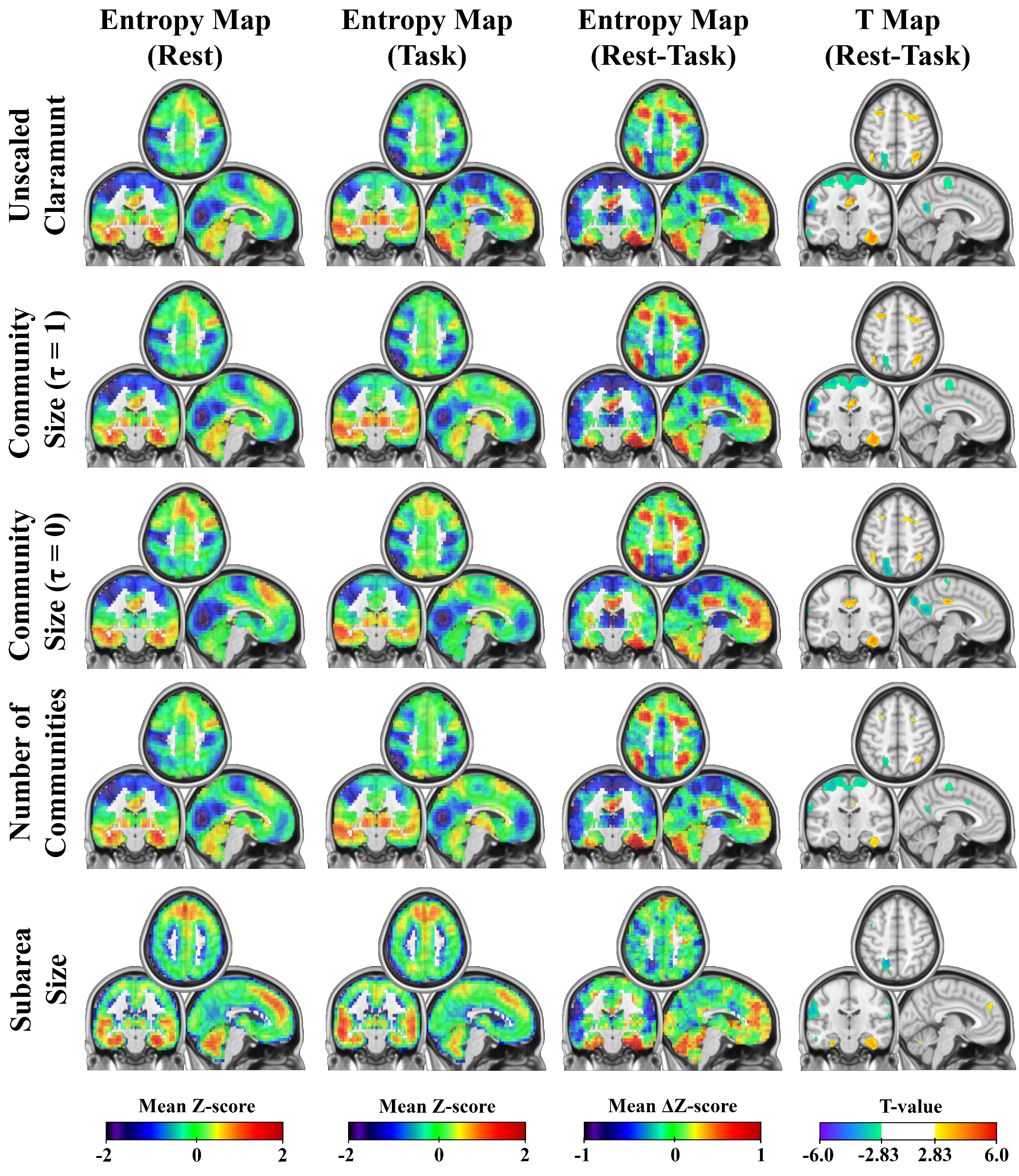

**Supplementary Figure 8.** Sample average entropy maps for resting-state (Rest, column 1) and 2-back (Task, column 2), the difference between tasks (column 3), and maps of statistically significant (without multiple comparisons correction) t-values for paired comparisons of resting-state and 2-back (column 4). Note that color scales change between the entropy maps (columns 1 and 2) and the difference maps (column 3). Each row corresponds to a different set of scaling parameters in comparison to Claramunt’s entropy measure without scaling (row 1).

Shown slices have MNI coordinates X = -5 , Y = -19, Z = 50

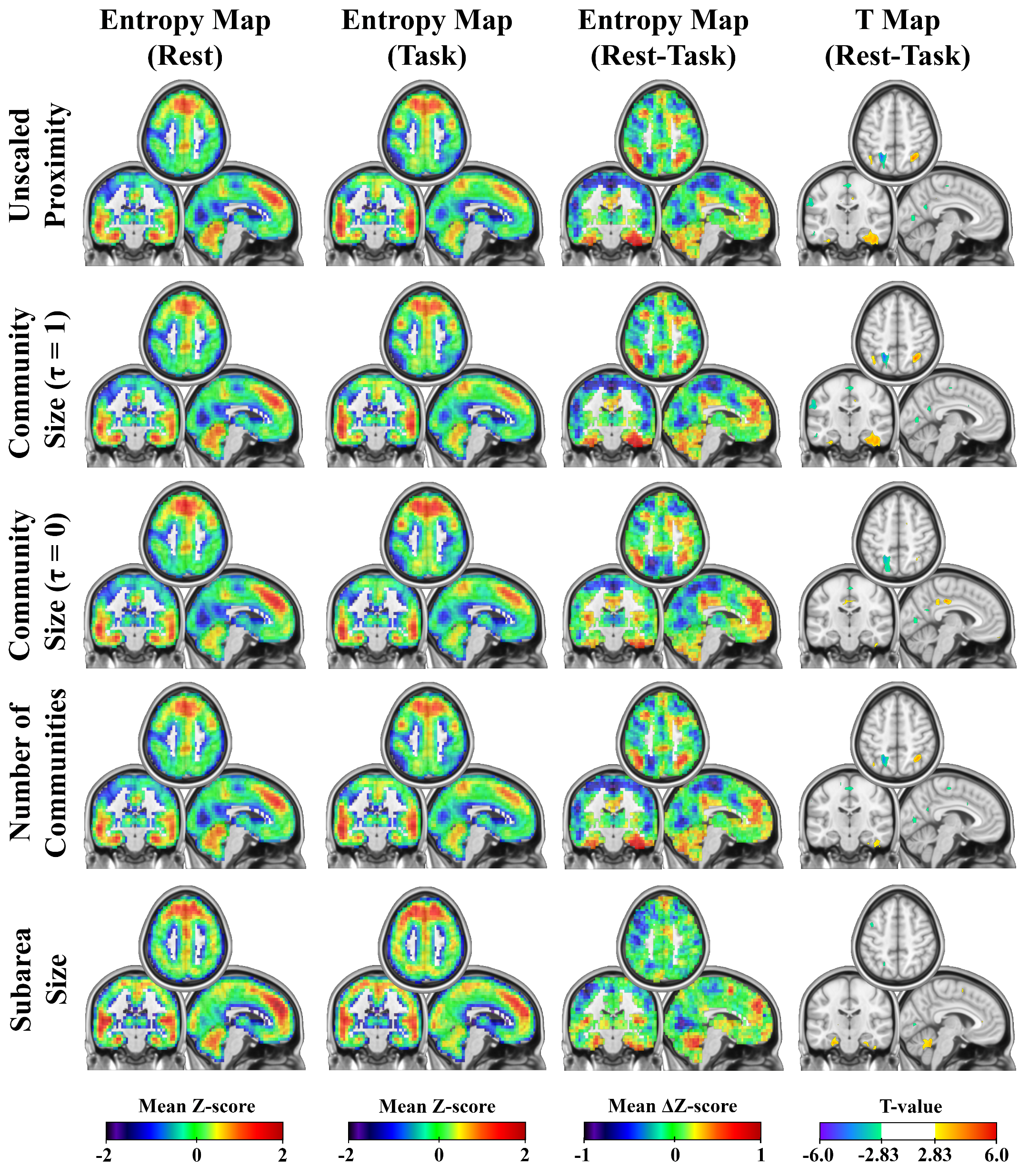

**Supplementary Figure 9.** Sample average entropy maps for resting-state (Rest, column 1) and 2-back (Task, column 2), the difference between tasks (column 3), and maps of statistically significant (without multiple comparisons correction) t-values for paired comparisons of resting-state and 2-back (column 4). Note that color scales change between the entropy maps (columns 1 and 2) and the difference maps (column 3). Each row corresponds to a different set of scaling parameters in comparison to the Proximity entropy measure without scaling (row 1).

Shown slices have MNI coordinates X = -5 , Y = -19, Z = 50

**Supplementary Table 1. Rest versus 2-back cluster-level results – Shannon’s Entropy**

| **Region** | **Peak MNI Coordinate**  **(x y z)** | **Cluster Extent** | **Cluster *p*-value** | **Peak *T*-value** |
| --- | --- | --- | --- | --- |
| Rest Entropy > Task Entropy | | | | |
| R. Middle and Superior Frontal Gyri | 34 6 53 | 112 | 0.010 | 6.72 |
| Midcingulate Cortex | 10 -18 33 | 66 | 0.039 | 5.87 |
| L. Middle and Superior Frontal Gyri | -30 14 48 | 60 | 0.048 | 5.81 |
| L. Cerebellar Crus I (extending into Inferior Temporal Gyrus) | -46 -54 -37 | 233 | 0.001 | 5.77 |
| L. Intraparietal Lobule and Angular Gyrus | -38 -46 43 | 85 | 0.022 | 5.38 |
| R. Superior Parietal Lobule | 38 -58 58 | 130 | 0.006 | 5.33 |
| R. Inferior Temporal and Fusiform Gyri (extending into Cerebellar Crus I) | 42 -42 -17 | 226 | 0.001 | 5.30 |
| R. Orbital Gyri | 22 34 -12 | 98 | 0.015 | 4.84 |
| Rest Entropy < Task Entropy | | | | |
| R. Inferior Frontal (triangle) and Middle Frontal Gyri | 46 38 13 | 79 | 0.026 | 6.70 |
| L. Supramarginal Gyrus (extending into Superior and Middle Temporal Gyri) | -62 -30 28 | 200 | 0.001 | 6.25 |
| L. and R. Pre- and Post-Central gyri | 18 -22 73 | 337 | <0.001 | 5.30 |
| R. Supramarginal Gyrus | 62 -38 28 | 81 | 0.024 | 5.29 |
| L. Middle Occipital Gyrus (extending to L. Fusiform gyrus) | -42 -86 23 | 129 | 0.006 | 5.15 |

**Supplementary Table 2. Rest versus 2-back cluster-level results – O’Neill’s Entropy**

| **Region** | **Peak MNI Coordinate**  **(x y z)** | **Cluster Extent** | **Cluster *p*-value** | **Peak *T*-value** |
| --- | --- | --- | --- | --- |
| Rest Entropy > Task Entropy | | | | |
| L. Cerebellar Crus I (extending into Inferior Temporal Gyrus) | -46 -50 -37 | 284 | <0.001 | 6.24 |
| R. Inferior Temporal and Fusiform Gyri (extending into Cerebellar Crus I) | 42 -42 -17 | 254 | 0.001 | 5.86 |
| R. Middle and Superior Frontal Gyri | 34 6 58 | 87 | 0.025 | 5.61 |
| R. Superior Parietal Lobule | 38 -58 58 | 102 | 0.017 | 5.15 |
| R. Orbital Gyri | 26 22 -17 | 116 | 0.012 | 4.86 |
| R. Parahippocampal Gyrus | 22 -34 -7 | 71 | 0.040 | 4.61 |
| Rest Entropy < Task Entropy | | | | |
| L. Supramarginal Gyrus (extending into Superior and Middle Temporal Gyri) | -62 -30 28 | 217 | 0.001 | 6.41 |
| R. Inferior Frontal (triangle) and Middle Frontal Gyri | 46 38 13 | 71 | 0.040 | 6.24 |
| L. Middle Occipital Gyrus (extending to L. Fusiform gyrus) | -42 -86 23 | 138 | 0.007 | 6.05 |
| L. and R. Precuneus | 2 -58 23 | 80 | 0.031 | 5.84 |
| R. Supramarginal Gyrus | 58 -38 38 | 84 | 0.027 | 5.33 |
| L. and R. Pre- and Post-Central gyri | 18 -22 73 | 267 | <0.001 | 4.71 |

**Supplementary Table 3. Rest versus 2-back cluster-level results – Claramunt’s Entropy**

| **Region** | **Peak MNI Coordinate**  **(x y z)** | **Cluster Extent** | **Cluster *p*-value** | **Peak *T*-value** |
| --- | --- | --- | --- | --- |
| Rest Entropy > Task Entropy | | | | |
| R. Inferior Temporal and Fusiform Gyri (extending into Cerebellar Crus I) | 42 -42 -22 | 293 | <0.001 | 6.86 |
| Midcingulate Cortex | 10 -14 38 | 48 | 0.028 | 5.99 |
| L. Cerebellar Crus I (extending into Inferior Temporal Gyrus) | -54 -58 -32 | 203 | <0.001 | 5.90 |
| R. Superior Parietal Lobule and Angular Gyrus | 30 -66 58 | 90 | 0.004 | 5.45 |
| L. Intraparietal Lobule and Angular Gyrus | -38 -62 43 | 55 | 0.020 | 5.43 |
| R. Orbital Gyri | 30 34 -12 | 79 | 0.007 | 5.38 |
| R. Middle and Superior Frontal Gyri | 30 6 48 | 48 | 0.028 | 4.45 |
| R. Parahippocampal Gyrus | 14 -30 -7 | 49 | 0.027 | 4.20 |
| Rest Entropy < Task Entropy | | | | |
| L. Supramarginal Gyrus (extending into Superior and Middle Temporal and Pre- and Post-Central Gyri) | -62 -30 28 | 462 | <0.001 | 6.93 |
| R. Supramarginal Gyrus | 66 -38 38 | 94 | 0.004 | 5.65 |
| L. Inferior Temporal Gyrus (extending to Fusiform and Middle Occipital Gyri) | -46 -58 -7 | 146 | 0.001 | 5.45 |
| R. Inferior Frontal (triangle) and Middle Frontal Gyri | 42 62 3 | 61 | 0.015 | 5.18 |
| L. Superior Parietal Lobule (extending to Superior Occipital Lobule) | -18 -66 38 | 77 | 0.007 | 5.05 |
| L. and R. Precuneus | 2 -58 23 | 46 | 0.031 | 4.90 |

**Supplementary Table 4. Rest versus 2-back cluster-level results – Proximity Entropy**

| **Region** | **Peak MNI Coordinate**  **(x y z)** | **Cluster Extent** | **Cluster *p*-value** | **Peak *T*-value** |
| --- | --- | --- | --- | --- |
| Rest Entropy > Task Entropy | | | | |
| R. Cerebellar Crus I (Extending throughout R. Cerebellum | 54 -54 -32 | 127 | <0.001 | 5.74 |
| R. Temporal Pole | 46 18 -27 | 26 | 0.038 | 5.61 |
| L. Cerebellar Crus I (extending into Fusiform Gyrus) | -30 -26 -32 | 81 | 0.001 | 5.23 |
| R. Superior Parietal Lobule and Angular Gyrus | 30 -58 48 | 49 | 0.007 | 4.96 |
| L. Intraparietal Lobule and Angular Gyrus | -38 -62 43 | 40 | 0.013 | 4.74 |
| R. Parahippocampal Gyrus (extending to Inferior Temporal and Fusiform Gyri) | 38 -14 -22 | 95 | <0.001 | 4.61 |
| L. Cerebellar Lobule VIII | -14 -58 -57 | 23 | 0.049 | 3.66 |
| Rest Entropy < Task Entropy | | | | |
| L. Supramarginal Gyrus | -58 -30 28 | 65 | 0.002 | 6.12 |
| L. Fusiform Gyrus (extending to Inferior Temporal Gyrus) | -34 -62 -12 | 100 | <0.001 | 5.55 |
| L. Superior Parietal Lobule (extending to Superior Occipital Lobule) | -22 -54 48 | 66 | 0.002 | 5.51 |
| L. Middle Occipital Lobule | -42 -82 23 | 61 | 0.003 | 5.28 |
| R. Supramarginal Gyrus | 66 -34 38 | 33 | 0.021 | 4.97 |
| R. Middle Frontal Gyrus | 42 58 3 | 28 | 0.032 | 4.93 |
| R. Inferior Temporal Gyrus (extending to Middle Occipital Lobule) | 54 -66 -7 | 88 | 0.001 | 4.77 |
